# GAPIT Version 3: Boosting Power and Accuracy for Genomic Association and Prediction

**DOI:** 10.1101/2020.11.29.403170

**Authors:** Jiabo Wang, Zhiwu Zhang

## Abstract

Genome-Wide Association Study (GWAS) and Genomic Prediction/Selection (GP/GS) are the two essential enterprises in genomic research. Due to the great magnitude and complexity of genomic data, analytical methods and their associated software packages are frequently advanced. GAPIT is a widely used Genomic Association and Prediction Integrated Tool. The first version was released to the public in 2012 with the implementation of the general linear model (GLM), mixed linear model (MLM), compressed MLM, and genomic Best Linear Unbiased Prediction (gBLUP). The second version was released in 2016 with several new implementations, including Enriched Compressed MLM and Settlement of mixed linear models Under Progressively Exclusive Relationship (SUPER). All the GWAS methods are based on the single locus test. For the first time, in the current release of GAPIT, version 3 implemented three multiple loci test methods, including Multiple Loci Mixed Model (MLMM), Fixed and random model Circulating Probability Unification (FarmCPU), and Bayesian-information and Linkage-disequilibrium Iteratively Nested Keyway (BLINK). Additionally, two GP/GS methods were implemented based on Compressed MLM, named compressed BLUP, and SUPER, named SUPER BLUP. These new implementations not only boost statistical power for GWAS and prediction accuracy for GP/GS, but also improve computing speed and increase the capacity to analyze big genomic data. Here, we document the current upgrade of GAPIT by describing the selection of the recently developed methods, their implementation, and potential impact. All documents, including source code, user manual, demo data, and tutorials, are freely available at the GAPIT website (http://zzlab.net/GAPIT).

## Introduction

Computer software is essential tool for genomic research. Genome-wide association studies (GWAS) and genomic prediction are the two essential enterprises for genomic research. For a particular trait of interest, GWAS focuses on finding genetic loci associated with the causal genes and estimating their effects. Genomic prediction, known as genomic selection (GS) in the fields of animal and plant breeding, focuses on the direct prediction of phenotypes by estimating the total genetic merit underlying the phenotypes [1]. The estimated genetic merit is also known as the estimated breeding value (EBV) for animal and plant breeding. In the long term, the assessment of all genetic loci underlying a trait may eventually lead to highly accurate EBV predictions. In the short term, methods have been developed to derive EBV even without identifying those associated genetic loci. Consequently, some statistical methods are shared between GWAS and GS, and some methods are specific to each. Accordingly, the software packages are also characterized into GWAS-specific, GS-specific, or packages that perform both.

For GWAS, many statistical methods and software packages have been developed to improve computational efficiency, statistical power, and control of false positives. The most computational efficient method is the General Linear Model (GLM), which can fit population structure or principal components as fixed effects to reduce the false positives caused by population stratification[2,3]. To account for the relationships among individuals within sub-populations, kinship among individuals was introduced through the mixed linear model (MLM) by using genetic markers covered the entire genome[4]. This strategy served to further control false positives. To reduce the computational burden of MLM, many algorithms have been developed, including Efficient Mixed Model Association (EMMA)[5], EMMA eXpredited (EMMAx), Population Parameter Previously Determined (P3D)[6,7], factored spectrally transformed linear mixed models (FaST-LMM) [8], and GRAMMAR-Gamma[9]. These methods improve computing efficiency of MLM, but their statistical power remain the same as MLM.

Enhancement of MLM have also been introduced to improve statistical power. To reduce the confounding between kinship and testing markers, individuals in the MLM are replaced with their corresponding groups in the compressed MLM (CMLM), which also improves computing efficiency[7]. Refer to the cluster method to fit such relationship between individuals, the enriched CMLM (ECMLM) was developed to further improve statistical power[10]. Instead of using all markers to derive kinship among individuals across traits of interest, selection of the markers according traits of interest can improve statistical power. One of such methods is the Settlement of MLM Under Progressively Exclusive Relationship (SUPER)[11]. SUPER contains three steps. The first step was the same as other models such as GLM or MLM to have a initiate assessment of the marker effects. In the second step, kinship is optimized using maximum likelihood in a mixed model with kinship derived from the selected markers based on their effects and relationship on linkage disequilibrium. In the third step, markers are tested again one at a time as final output with kinship derived from the selected markers except the ones that are in linkage disequilibrium with the testing markers.

Same as the extension of single-marker tests using GLM to stepwise regression (e.g. GLMSelect Procedure in SAS)[12,13], single-locus tests using MLM were also extended to multiple loci tests, named multiple loci mixed linear model (MLMM) [14]The most significant maker is fitted as a covariate in the stepwise fashion. The iteration stops when variance associated with the kinship goes to zero, followed by a backward stepwise regression to eliminate the non-significant covariate markers. In MLMM, both covariate markers and kinship are fitted in the same MLM. This model was separated into two models which are iterated back and forth. One model is MLM which contains the random effect associated with kinship and covariates such as population structure, but not the associate markers. The associated markers are optimized to derive the kinship using maximum likelihood. The other model is a GLM containing a testing mark and covariates such as population structure. The method was named as Fixed and random model Circulating Probability Unification (FarmCPU) [15]. Because a marker test in GLM does not involve kinship, FarmCPU is not only faster but gives higher statistical power than MLMM. The MLM in FarmCPU was further replaced with GLM to speed up in the new method named the Bayesian-information and Linkage-disequilibrium Iteratively Nested Keyway (BLINK) [16]. The maximum likelihood method in MLM was replaced by the Bayesian-information content. BLINK eliminates the restriction assuming that causal genes are evenly distributed across the genome by SUPER and FarmCPU method, consequently boosting statistical power.

For genomic prediction/selection, the earliest effort can be traced to the use of marker-based kinship in the Best Linear Unbiased Prediction (BLUP) method, currently known as genomic BLUP or gBLUP [17–19]. The method uses all markers covering the whole genome to define the kinship among individuals to estimate their EBV. A different strategy is to estimate the effects of all markers and sum them together to predict individuals’ total genetic effects [20]. To avoid the overfitting problem in the fixed-effect model, these markers are fitted as random effects simultaneously. A variety of restrictions and assumptions are applied to these random effects and their prior distributions under the Bayesian theorem. Different methods were named according to different priors, such as Bayes A, B, Cpi, and LASSO [20]. The case assuming the effect of all markers have the same distribution with constant prior variance is equivalent to Ridge Regression [18,21].

Many of the software package developments accompanied GWAS and GS method developments so that the methods and the software were given the same name, such as EMMA[5], EMMAx[22], FaST-LMM[8], FarmCPU [15], and BLINK[16]. Often, to compare different statistical methods, users must learn how to use the various software packages. To reduce the multiple steep learning curves for users, some packages were developed with more than one statistical method. These packages include PLINK with GLM and logistic regression [23]; TASSEL [24] with GLM and MLM; rrBLUP with ridge regression and gBLUP [25]; and BGLR with ridge regression, gBLUP, and Bayesian methods [26]. Also, some packages have implemented methods for both GWAS and GS so that users can use one software package to conduct both analyses. One example is Genome Association and Prediction Integrated Tool (GAPIT). GAPIT was initiated with GLM, MLM, EMMAx/P3D, CMLM, and gBLUP in version 1 [27] and enriched with ECMLM, FaST-LMM, and SUPER in version 2[28].

Furthermore, with such a variety of available methods, researchers feel extremely overwhelmed when trying to choose the best method to analyze their particular data. This dilemma is especially true when only a subset of these methods has been compared under conditions less relevant to a researcher’s specific study conditions. For example, simulation studies have demonstrated that FarmCPU is superior to MLMM for GWAS [15]; however, no comparisons have been conducted between SUPER and FarmCPU or between SUPER and MLMM. Similarly, for GS, gBLUP, SUPER BLUP (sBLUP), and Compression BLUP (cBLUP) have been compared with Bayesian LASSO [1]. Thus, software packages with features that allow researchers to conduct comparisons for model selection—especially under the conditions relevant to their studies—are critically needed.

Moreover, because the results of existing software packages are displayed as static output, researchers often find that extracting relevant information is challenging. For example, users must spend additional effort searching through file outputs to obtain the estimated effect and minor allele frequency (MAF) for a particular marker observed on Manhattan and QQ plots. Yet, this extra effort is necessary because these two factors are essential to infer the causes of association. For 3-D plots of population structure, users are unable to identify properties that are currently hidden by the angles determined by the software. The capability of angle adjustment would largely resolve this issue. Therefore, researchers are also in critical need of an interactive, dynamic output display system that allows flexibility, easy extraction of relevant information.

To address these critical needs, we continuously strive to upgrade GAPIT software by adding state-of-the-art GWAS and GS methods as they become available. Herein, we report our most recent efforts to upgrade GAPIT to version 3 (GAPIT3) by implementing MLMM, FarmCPU and BLINK [14–16] for GWAS, and sBLUP and cBLUP for GS[1]. We also added features that allow users to interact with both the analytical methods and displayed outputs for comparison and interpretation. Users’ prior knowledge can now be used to enhance method selection and unfold the discoveries hidden by static outputs.

## Methods

### Architecture of GAPIT version 3

To implement three multiple-locus GWAS methods (MLMM, FarmCPU, and BLINK) and two new methods of GS (cBLUP and sBLUP), we redesigned GAPIT with a new architecture to easily incorporates an external software package. In order of execution, GAPIT is compartmentalized into five modules: 1) Data and Parameters (DP); 2) Quality Control (QC); 3) Intermediate Components (IC); 4) Sufficient Statistics (SS); and 5) Interpretation and Diagnoses (ID). Any of these modules are optional and can be skipped. However, GAPIT3 does not allow modules to be executed in reverse order (**Figure 2**).

The DP module contains functions to interpret input data, input parameters, genotype format transformation, missing genotype imputation, and phenotype simulations. The types of input data and their labels are the same as previous versions of GAPIT, including phenotype data (Y); genotype data in either Hapmap format (G), or numeric data format (GD) with genetic map (GM); covariate variables (CV), and kinship (K). The input parameters include those from previous GAPIT versions plus the parameters for the new GWAS and GS methods and the enrichments associated with the other four modules. Two genetic models, additive and dominant, are available to transform genotypes in HapMap format into numeric format. Under the additive model, homozygous genotypes with recessive allele combinations are coded 0, homozygous genotypes with dominant allele combinations are coded 2, and heterozygous genotypes are coded 1. Under the dominant model, both types of homozygous genotypes are coded 0 and heterozygous genotypes are coded 1. When genotype, heritability, and number of QTNs are provided without phenotype data, GAPIT3 will conduct a phenotype simulation from the genotype data.

By default, GAPIT3 assumes users provide quality data and does not perform data quality control. When the quality control option is turned on, GAPIT will conduct quality control on imputing missing genotypes, filtering markers by MAF, sorting individuals in phenotype and genotype data, and matching the phenotype and genotype data together. GAPIT provides multiple options for genotype imputation, including major homozygous genotypes and heterozygous genotypes.

In the IC module, GAPIT provides comprehensive functions to generate intermediate graphs and reports, including phenotype distribution, MAF distribution, heterozygosity distribution, marker density, LD decay, principal components, and kinship. These reports and graphs help users to diagnosis and identify problems with the input data for quality control. For example, an associated marker should be further investigated if it has low MAF.

The SS module contains multiple adapters that generate sufficient statistics for existing methods in the previous versions of GAPIT and new external methods. The sufficient statistics are the P values for GWAS and predicted phenotypes for GS. The methods in the previous versions include GLM, MLM, CMLM, ECMLM, SUPER, and gBLUP. The new adapters developed in GAPIT3 include MLMM, FarmCPU, BLINK, cBLUP, and sBLUP.

The ID module contains the static reports developed in previous GAPIT versions and the new interactive reports generated in GAPIT3. The interactive reports include the rotational three-dimensional plot of the first three principal components, display of marker information on Manhattan plots and QQ plots, and individual information on the phenotype plots (predicted vs. the observed). The marker information includes maker name, chromosome, position, MAF, and effect estimate. The individual information consists of the individual name and the values for predicted and observed phenotypes.

### Implementation of MLMM and FarmCPU

Both MLMM and FarmCPU have source code available on their websites. These source codes were directly integrated into the GAPIT source code, so users are only required to install GAPIT3, not all three packages. We also added the input parameters specific to MLMM and FarmCPU into the input parameter list of GAPIT3. These two software packages share a similar input and output data format for phenotypes, genotypes, covariate variables, and P values. GAPIT currently does not support some formats for genotype data, including objects with bigmemory and biganalytics. Consequently, the data scale that can be processed by FarmCPU is larger than GAPIT for using FarmCPU GWAS method.

Integrating MLMM and FarmCPU source code into GAPIT source code lowers the risk of breaking the linkage between GAPIT and these two software packages when they release updates. The disadvantage is that MLMM and FarmCPU source codes remain static in GAPIT. The GAPIT team periodically checks for updates of these two packages and correspondingly updates the GAPIT source code.

### Implementation of Blink R and C versions

BLINK R version was released as an executable R package on GitHub. GAPIT accesses BLINK R as an independent package. The BLINK C version was released as an executable C package on GitHub. To access BLINK C, GAPIT needs the executable program in the working directory. To avoid the potential risk of breaking the linkage between GAPIT and BLINK, the GAPIT team maintains a close connection with the BLINK team for updates. BLINK C conducts analyses on binary files for genotypes. The binary files not only make BLINK C faster, but also provide the capacity to process big data with limited memory. Running BLINK C through GAPIT requires nonbinary files first, then BLINK C is used to convert them to binary. For big data, we recommend directly accessing BLINK C to obtain P values and using the GAPIT ID module to interpret and diagnosis the results.

### Implementation of cBLUP and sBLUP

The compressed BLUP (cBLUP) and SUPER BLUP (sBLUP) were developed from the corresponding GWAS methods: compressed MLM (CMLM) and SUPER. Because CMLM and SUPER were already implemented in GAPIT versions 1 and 2, respectively, implementation of cBLUP and sBLUP was more straightforward than other implementations. For cBLUP, the solutions of the random group effects in CMLM are used as the genomic estimated breeding values for the corresponding individuals. For sBLUP, the calculation is even easier than the SUPER GWAS method. For the SUPER GWAS method, a complementary kinship is used for a testing SNP that is in linkage disequilibrium with some of the associated SNPs. For sBLUP, all associated markers are used to derive the kinship and subsequently to predict the breeding values of individuals. No operation for the complementary process is necessary.

### Implementation of interactive reports

Two types of interactive reports are included in the current GAPIT3. First, users can now interact with Manhattan plots, QQ plots, and scatter plots of predicted vs. observed phenotypes to extract information about markers and individuals. For example, by moving the cursor or pointing device over a data point, users can find names and positions of markers or names and phenotypes of individuals. An R package plotly was used to store this type of information in the format of HTML files, which can be displayed by web browsers. Second, users can rotate graphs such as three-dimensional PC plots using a pointing device such as mouse or trackpad. The R packages (rgl and rglwidget) were jointly used to realize the functions.

### Proportion of variance explained

In GAPIT3, the proportion of total phenotypic variance explained by significantly associated markers is evaluated. A Bonferroni multiple test threshold is used to determine significance. The associated markers are fitted as random effects in a multiple random variable model. The model also include other fixed effects are used in the GWAS to select these associated markers. The multiple random variable model is analyzed using an R package, lme4, to estimate the variance of residuals and the variances of the associated markers. The proportions explained by the markers are calculated as their corresponding variances divided by the total variance, which is the sum of residual variance and the variance of the associated markers.

## Results

GAPIT is a widely used software package. GAPIT website received over 22,000 pageviews. The GAPIT forum on Google contains ~1600 posts covering ~400 topics regarding the usage, functions, bugs, and fixes. These posts were viewed ~3000 times by the GAPIT community between 2016 and 2019. During this period, GAPIT received 887 and 89 citations for version 1 and version 2 articles, respectively (**Figure S1 and S2**). The GAPIT3 project started after the 2016 publication of GAPIT version 2 (GAPIT2). Since then, we implemented three multiple locus methods for GWAS and two methods for GS (**Figure 1**). In addition, we enhanced the outputs of GAPIT to improve their quality and to help users more easily diagnose the data quality, compare analytical methods, and interpret the results.

**Figure 1.**
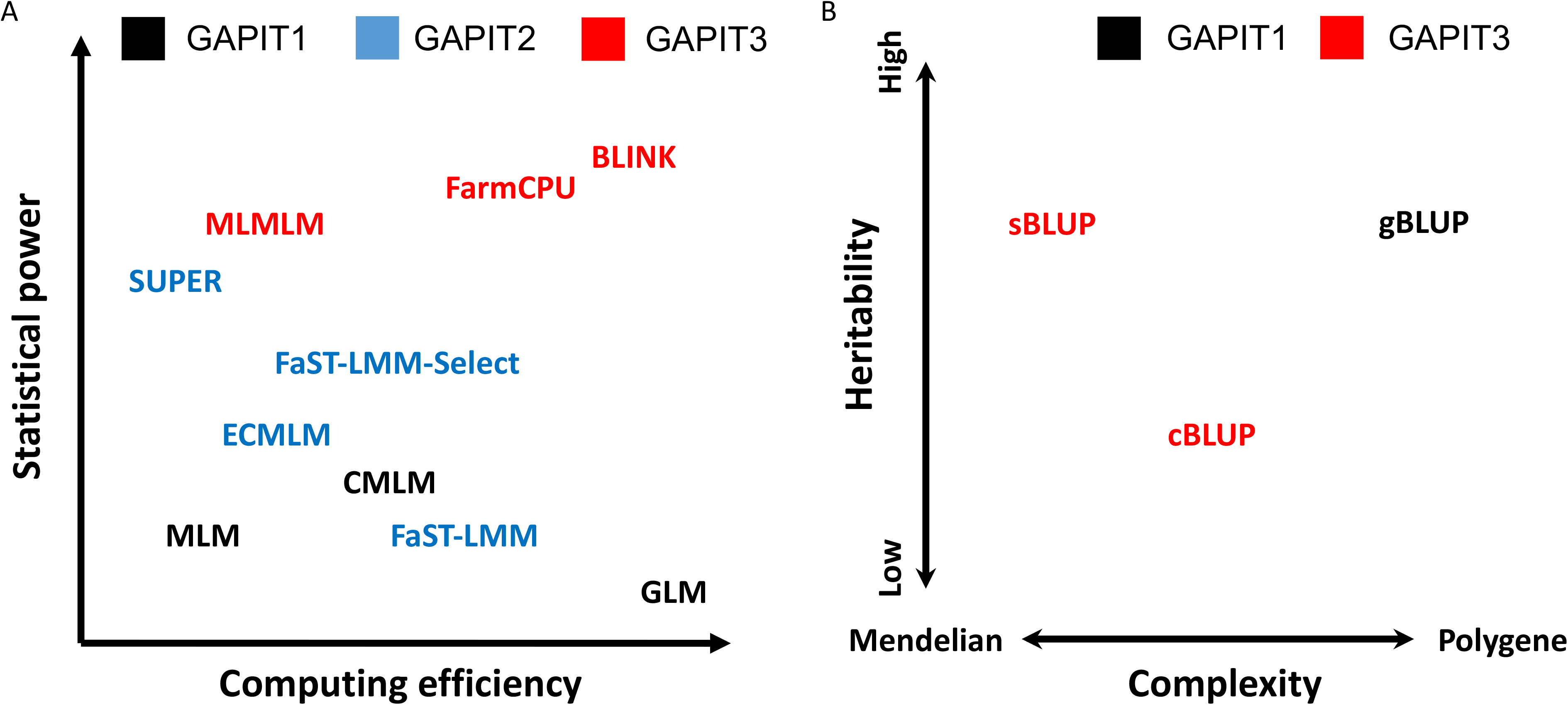
Statistical methods implemented in previous and current versions of GAPIT. The statistical methods are characterized by statistical power and computing efficiency (A) for genome-wide association study (GWAS) and by genetic architecture of targeting traits for Genomic Selection (GS) with respect to heritability and complexity (B). The GWAS methods include General linear model (GLM), Mixed linear model (MLM), compressed MLM (CMLM), factored spectrally transformed linear mixed models (FaST-LMM), FaST-LMM-Select, enriched CMLM (ECMLM), and settlement of mixed linear models under progressively exclusive relationship (SUPER). The GS methods include the regular genomic Best Linear Unbiased Prediction (gBLUP), compressed BLUP (cBLUP), and SUPER BLUP (sBLUP). Methods in black text were the ones implemented in the initial version of GAPIT, methods in blue text were new in GAPIT2, and methods in red text are new in the current GAPIT3.

**Figure 2.**
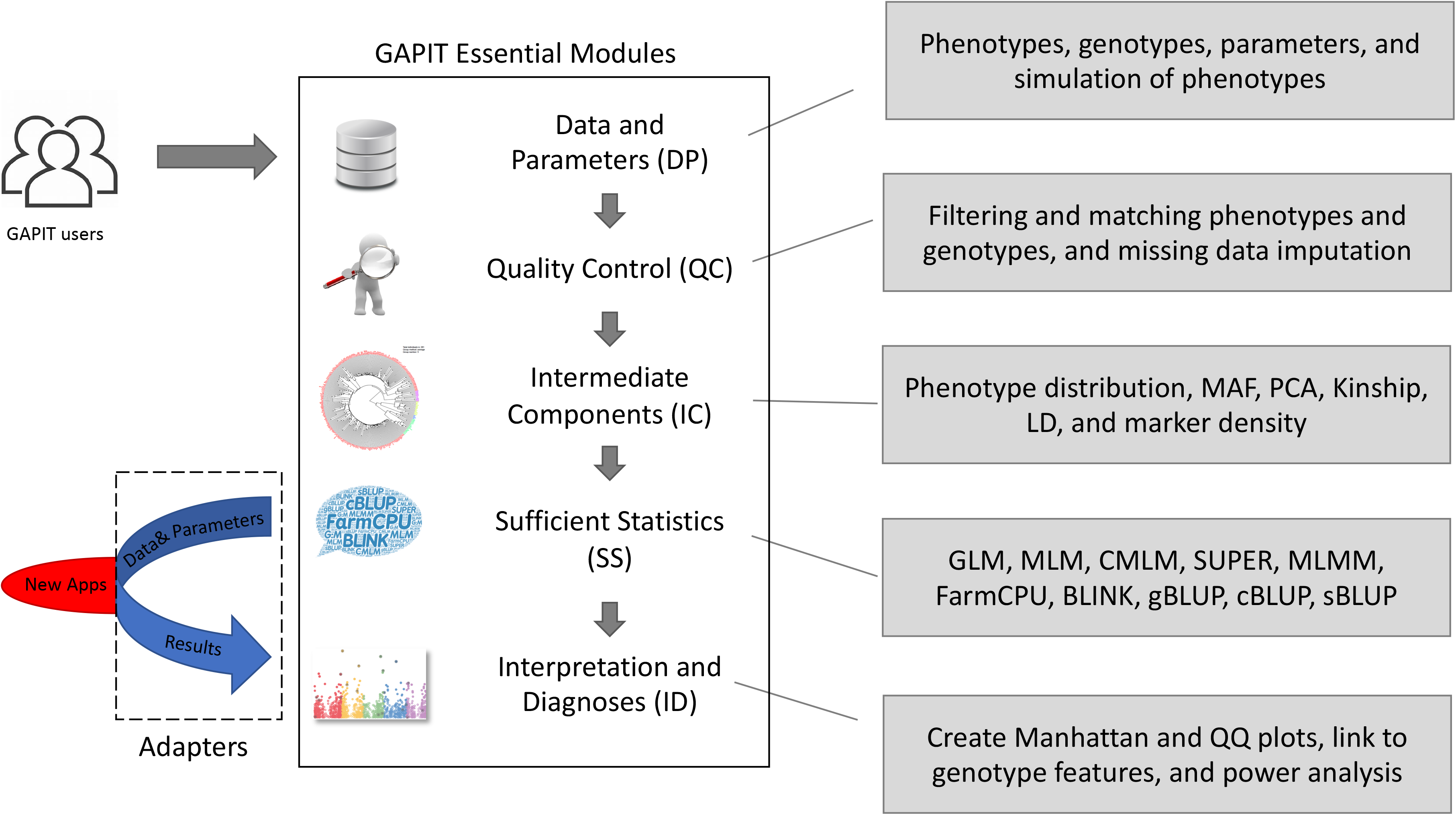
GAPIT essential modules and adapters to external packages. GAPIT version 3 was designed to have five sequential modules and multiple adapters that connect external software packages. The first module (DP) is responsible to process input data and parameters from users. The second module (QC) is responsible for quality control, including missing genotype imputation. The third module (IC) provides intermediate results, including Minor Allele Frequency (MAF), Principal Component Analysis (PCA), kinship, Linkage Disequilibrium (LD) analysis, and maker density distribution. The fourth module (SS) contains multiple adapters that convert input data into sufficient statistics, including maker effects, P values, and predicted phenotypes. The current adapters include General Linear Model (GLM), Mixed Linear Model (MLM), Compressed MLM (CMLM), SUPER (Settlement of MLM Under Progressively Exclusive Relationship), Multiple Locus Mixed Model (MLMM), FarmCPU (Fixed and random model Circulating Probability Unification), BLINK (Bayesian-information and Linkage-disequilibrium Iteratively Nested Keyway), genomic Best Linear Unbiased Prediction (gBLUP), Compressed BLUP, and SUPER BLUP (sBLUP). The fifth module provides the interpretation and diagnosis on the final results, included P values illustrated as Manhattan plots and QQ plots.

### Implementation of GWAS and GS methods

GAPIT version 1 (GAPIT1) was initiated with the single-locus test based on the CMLM, which clusters individuals into groups based on kinship. Because the CMLM is in a general format covering GLM and regular MLM, GAPIT can also conduct the MLM and the GLM. The MLM is equivalent to assigning each individual as its own group; the GLM is equivalent to assigning all individuals into one group. Consequently, CMLM is an optimization between MLM and GLM. The computation complexity of MLM is cubic to the number of individuals; thus, compression of individuals to groups not only improves statistical power, but also dramatically reduces computing time (**Figure 1A)**.

To improve the computing speed of MLM, GAPIT2 implemented FaST-LMM, which uses a set of markers to define kinship without performing the actual calculations. To further improve the statistical power of CMLM, the ECMLM was implemented to optimize the group kinship. Furthermore, two similar methods, SUPER and FaST-LMM-Select, were implemented in GAPIT2 to use a kinship that is complementary to testing markers.

All GWAS methods implemented in GAPIT1 and GAPIT2 are based on the single locus testing. The opposite approach, multiple loci tests, has received more attention since 2012, with the introduction of multiple loci mixed models (MLMM) using stepwise regression[14]. Through the use of iteration, two additional methods have been developed for multiple loci tests. The first method, Fixed and random model Circulating Probability Unification (FarmCPU); uses iteration between a fixed effect model and a random effect model. The second method, Bayesian-information and Linkage-disequilibrium Iteratively Nested Keyway (BLINK), uses iteration between two fixed-effect models. In GAPIT3, we implemented all of three of these multiple loci test methods (MLMM, FarmCPU, and BLINK). We simulated 100 traits and ran four methods (GLM and MLM are single locus methods, FarmCPU and Blink are multiple loci methods). The result of power against FDR and power against type I error were used to compare the performance differences between single locus and multiple loci (**Figure S6**).

For genomic prediction or selection, GAPIT1 and GAPIT2 implement gBLUP using MLM. This method works well for traits controlled by many genes, but not as well for traits controlled by a small number of genes. To overcome this difficulty, the updated GAPIT3 implements the sBLUP method which is superior to gBLUP for traits controlled by a small number of genes[1]. Both gBLUP and sBLUP have a disadvantage for traits with low heritability. Therefore, GAPIT3 implements the cBLUP method [1]which is superior to both gBLUP and sBLUP for traits with low heritability (**Figure 1B**).

For most GWAS methods, GAPIT3 executes both GWAS and GS by default. This default option can be changed by including the statement “SNP.test=F” to conduct GS only. For GWAS with MLM and FaST-LMM, gBLUP is used for GS. For CMLM and ECMLM, cBLUP is used for GS. For SUPER and FaST-LMM-Select, sBLUP is used for GS. The exceptions are GLM, MLMM, FarmCPU, and BLINK. When these methods are selected, only GWAS is executed.

The new GAPIT3 creates two types of Manhattan plots, the standard orthogonal type with x- and y-axes (**Figure S3A**), and a circular type (**Figure S3B**) which take less display space. The overlap in results between multiple methods is displayed as either solid or dashed vertical lines that will extend through the Manhattan plots for all methods (**Figure S3**). A solid vertical line indicates that the overlap of significant SNP is shared by more than two methods and a dashed vertical line indicates the overlap is between only two methods. When multiple traits are analyzed with a single method, the trait results are displayed in the same style as multiple methods. When both multiple methods and multiple traits are employed, the method plots are nested within the trait plots.

### Adaptation of existing GAPIT users

Users already familiar with GAPIT software have experienced no difficulty migrating to version 3. Experiences of using other related software packages also help to use GAPIT. GAPIT generated identical results for the same methods implemented in the separated packages (**Figure 3**). By default, GAPIT3 conducts GWAS using the BLINK method, which has the highest statistical power and computing efficiency among all methods implemented. Users can change the default to other methods by including a model statement. For example, to use the FarmCPU method, the user would include the statement “model = “FarmCPU “” to override the default. The model options include GLM, MLM, CMLM, ECMLM, FaST-LMM, FaST-LMM-Select, SUPER, MLMM, FarmCPU, and BLINK.

**Figure 3.**
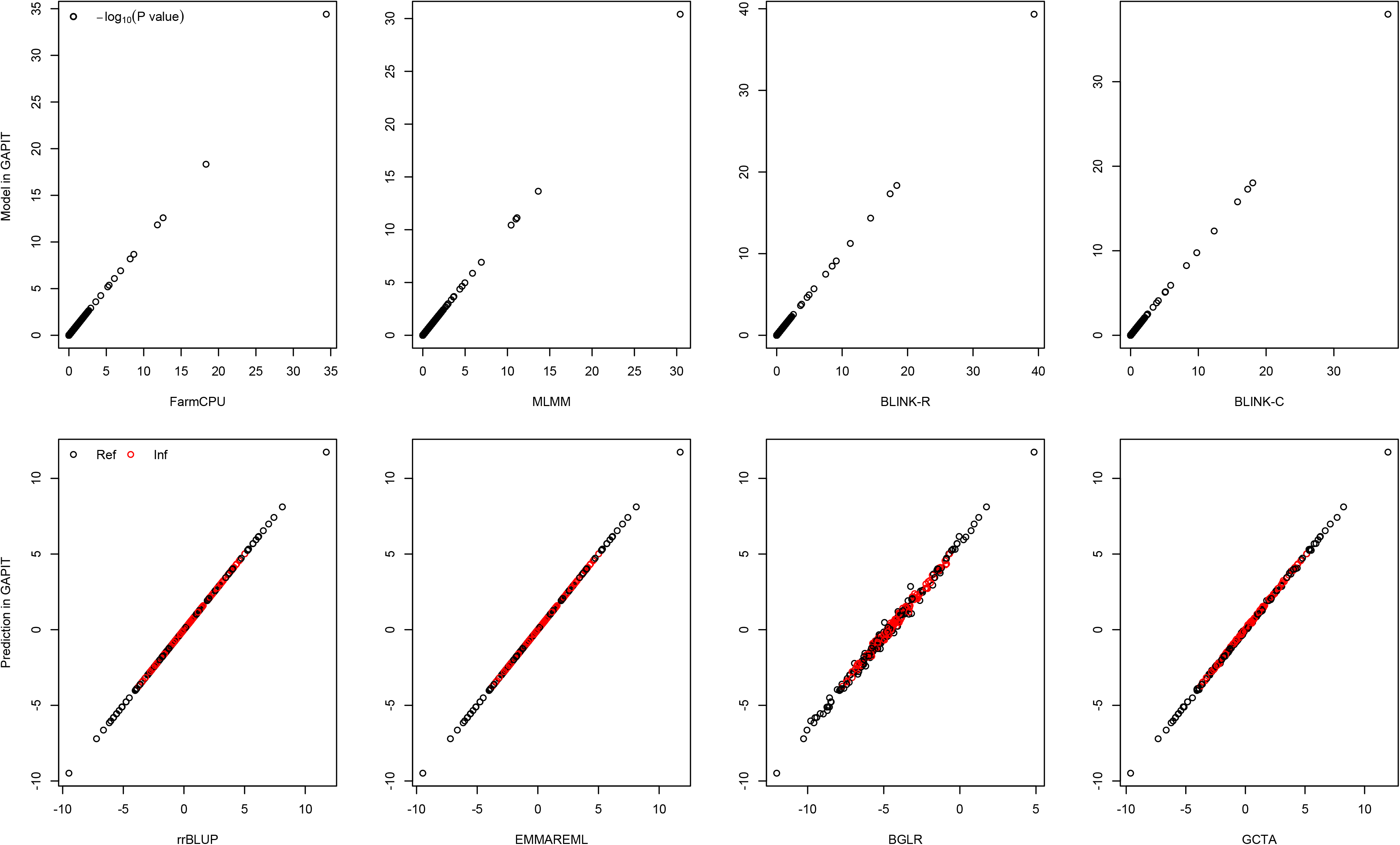
Comparison of P values and estimated breeding values using GAPIT and other software packages. The comparison was conducted on a trait simulated from the genotypes of 3093 SNPs on 281 maize lines. The simulated trait had 75% heritability with 20 QTNs. P values, displayed as -log10(P), are compared between GAPIT (vertical axis) and four software packages (horizontal axis) for genome-wide association studies that were run as standalone packages, including FarmCPU, MLMM, Blink R version, and BLINK C version. The estimated breeding values using GAPIT are compared with four software packages that were run as standalone packages, including rrBLUP, EMMAREML, BGLR, and GCTA. Identical results were obtained except breeding values using BGLR which involves random sampling to estimate variance components. The random sampling causes variation from run to run using BGLR.

GAPIT can also conduct GWAS and GS with multiple methods in a single analysis, allowing comparisons among methods for selection. For example, when the five methods (GLM, MLM, CMLM, FarmCPU, and BLINK) are used on maize flowering time in the demo data, inflation of p values and power of the analyses can be compared on the side-by-side Manhattan plots (**Figure S3**). All plots for the multiple methods show an interconnected vertical line that runs through chromosome 8. The results show that the GLM method identified association signals above the Bonferroni threshold (horizontal dashed red line in each plot). However, the association signals are inflated across the genome (the red dots on the QQ plots). BLINK method also identified two associated markers, including the marker close to a flowering time gene, VGT1 on chromosome 8. The QQ plot suggests that 99% of the markers have p values below the expected p values, which are indicated by the solid red line.

### Assessment of explained variance

GAPIT1 outputs the proportion of the regression sum of squares of testing markers to the total sum of squares as the estimate of variance explained by the markers. This approach is debatable because the sum of these proportions can exceed 100% when multiple markers are tested independently. In GAPIT2, this output was suppressed. However, we received substantial demands from GAPIT users for such output because some journals and reviewers require this information. To solve both of these problems, GAPIT3 conducts additional analyses using all associated markers as random effects. The proportion of variance of a marker over the total variance, including the residual variance, is reported as the proportion of total variance explained by the markers. This guarantees the sum of proportions of variance explained by the associated markers is below 100%. The non-associated markers are considered to contribute nothing to the total variance. The proportion of phenotypic variance explained by a marker is correlated with its minor allele frequency (MAF) and magnitude of marker effect. These relationships are demonstrated by scatter plots and a heatmap (**Figure 4**). The heat map indicates which markers explain a high proportion of the variance due to either a high MAF or a large magnitude of effect, or both.

**Figure 4.**
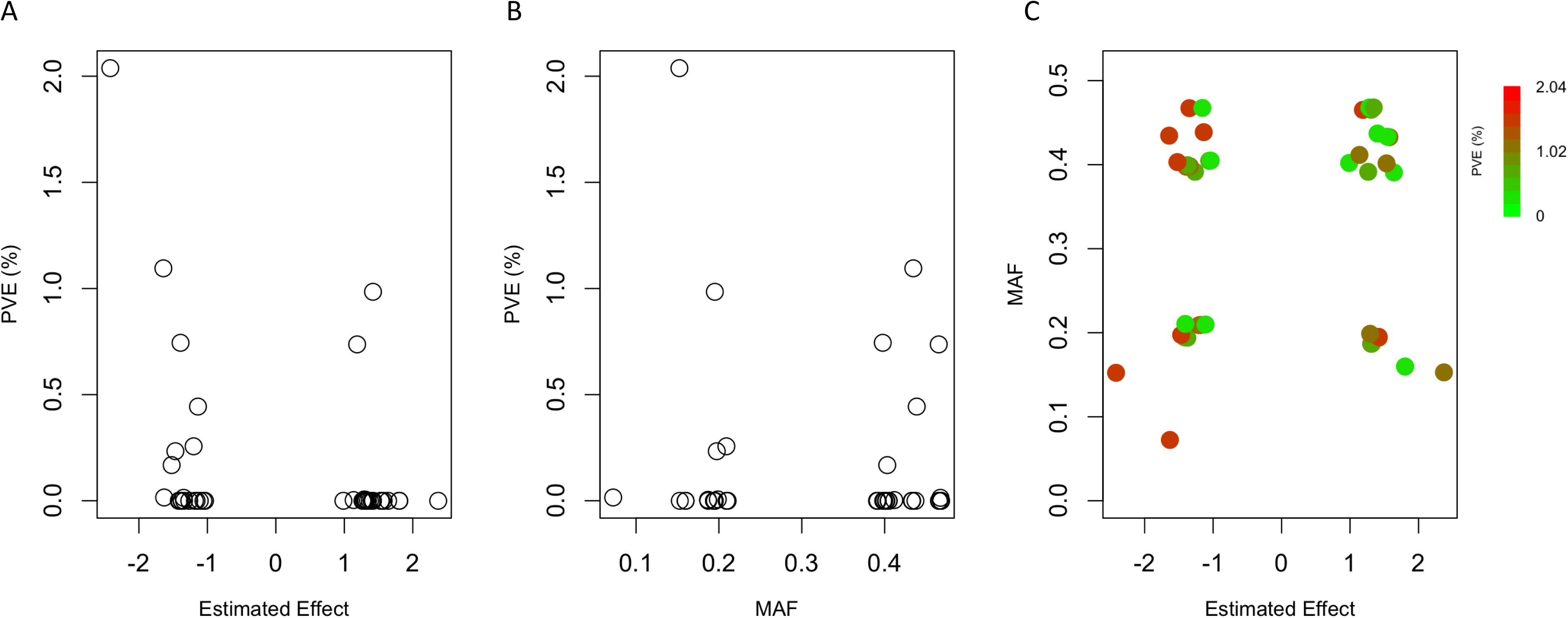
Phenotypic Variance Explained by Associated Markers. GAPIT 3 provides estimates of the proportion of phenotypic variance explained by associated markers. The proportion is a function of both magnitude of marker effects and minor allele frequency (MAF). Larger marker effects and larger MAF contribute to larger proportion of phenotypic variance explained. This relationship is demonstrated on a trait simulated from the mice genotypes of 12564 SNPs on 1440 individuals. The simulated trait had 75% heritability with 20 QTNs. Marker effects and MAF may go opposite direction. Some of markers have large magnitude, but explain little phenotypic variances due to low MAF (A). Similarly, markers with large MAFexplain little phenotypic variances due to small effect (B). Their joint impact is demonstrated by the heatmap (C). Markers explaining more variation are further away from the center where both MAF and marker effect are zeros.

### Enriched report output

When viewing the output graphics, such as Manhattan plots, QQ plots, and scatter plots of predicted vs. observed phenotypes, users are interested in the names and properties of markers and individuals. Finding this information usually requires computer programming to extract data from multiple resources, which includes searching files for P values, genotypes, estimated effects, and MAFs. With GAPIT3, in the interactive result all of information can be found by moving the cursor over the data point of interest (**Figure 5** and **S4**). For example, on the Manhattan and QQ plots, when the cursor moves over a data point, the marker information will be displayed. The Manhattan plot also contains a chromosome legend. Chromosomes can be hidden or displayed with different mouse clicking patterns. If a chromosome is clicked once, the plot will hide this chromosome; if clicked twice, the plot will hide all of the chromosomes besides chosen one. For the scatter plot of predicted vs. observed phenotypes, information about an individual is displayed when the cursor is moved over the associated data point of interest, including their names, observed, and predicted values.

**Figure 5.**
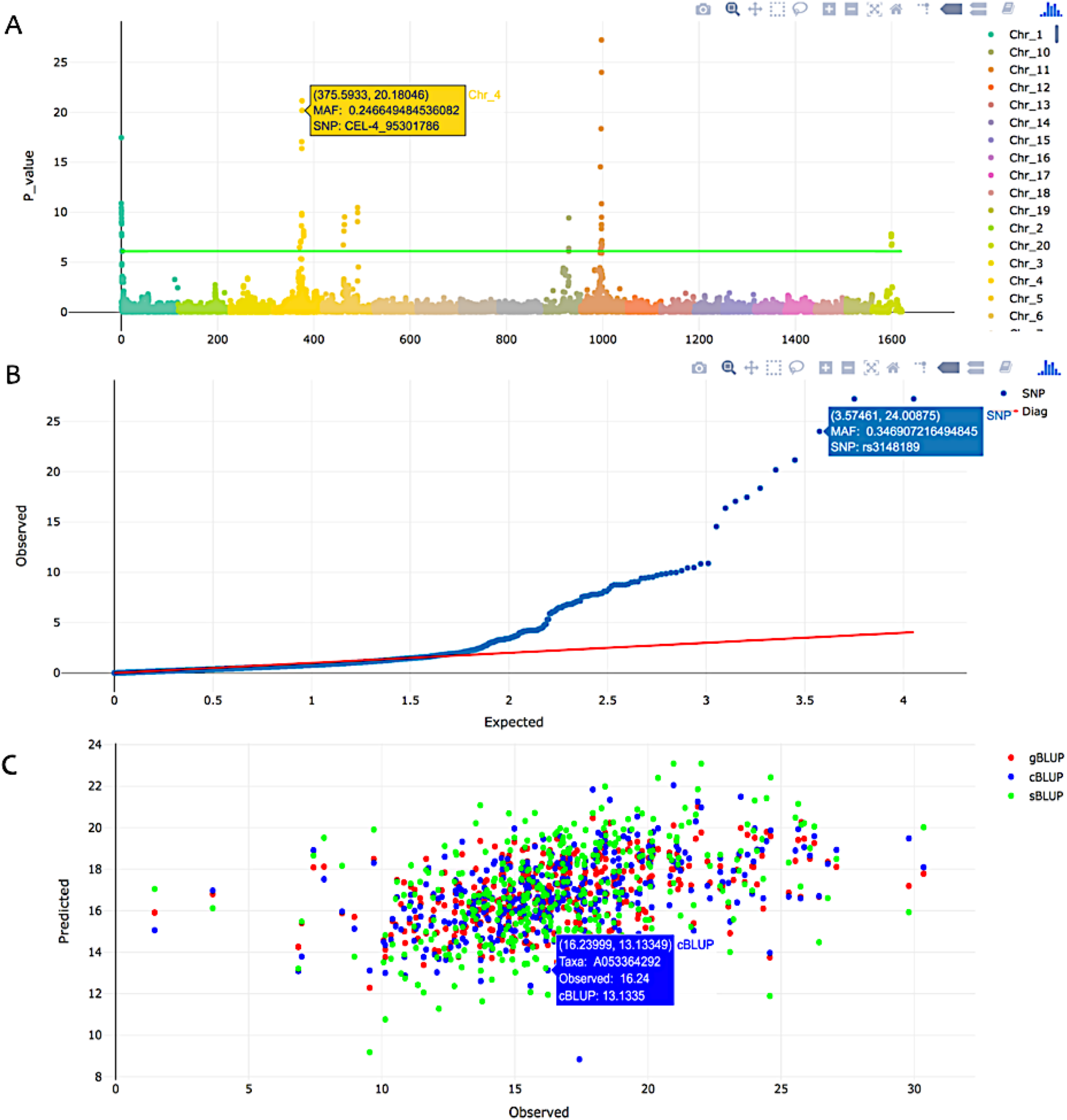
Interactive extraction of information for markers and individuals. GAPIT3 output two interactive html files to help user to extract information of markers on Manhattan plots (A) and QQ plots (B). The interactive plots are demonstrated on a trait simulated from the mice genotypes with 12564 SNPs on 1440 individuals. The simulated trait had 75% heritability with 20 QTNs. When cursor is moved over a dot, the marker information is displayed instantly, including name, P values, chromosome, position, and Minor Allele Frequency (MAF). Similarly, a html file is generated to display the predicted phenotypes against observed phenotypes (C). When cursor is moved over a dot, the individual information is displayed instantly, including name, predicted and observed phenotypic values. When multiple prediction methods are used, individuals are displayed as different colors for different methods, such as genomic Best Linear Unbiased Prediction (gBLUP), Compressed BLUP (cBLUP), and SUPER BLUP (sBLUP).

### Computing time

GAPIT3 newly implemented three multiple locus test methods (MLMM, FarmCPU, and BLINK) for GWAS and two methods (cBLUP and sBLUP) for genomic selection. All methods (GWAS and GS) have linear computing time to number of markers (**Figure 6AB**, and **S5**). However, they have mixed computing complexity to number of individuals. Most of them have computing time complexity that are cubic to number of individuals, including gBLUP and cBLUP for GS, and MLMM for GWAS. There are only two methods that have linear computing time to number of individuals: FarmCPU and BLINK (**Figure 6AB**). There is a minimal time increase for using MLMM. FarmCPU and BLINK packages within GAPIT from using them separately. There are two versions for BLINK methods: C version and R version. Literature demonstrated that the C version was much faster than the R version when they were operated as standard alone. When they were executed within GAPIT, the situation was reversed. This was because that GAPIT use the input and output directly for the R version. When GAPIT execute C version, the input and output data have to be transformed between memory and disk (**Figure 6AB**). For execution of gBLUP, GCTA was vigorous at all conditions to other packages, including BGLR, EMMREML, GAPIT and rrBLUP. All of these packages had linear computing time to number of markers, and nonlinear time to number of individuals. Their order changed depending number of individuals due to different setting cost. With number of markers duplicated four times and number of individuals duplicated at multiple levels (12, 20, and 28 fold), the computing show nonlinear relationship to number of individuals, except the GCTA package (**Figure 6C**). For small number of individuals (1124), BGLR was the slowest. When number of individuals was increased to three-fold (1124×3), rrBLUP became the slowest (**Figure 6DE**).. Therefore, GCTA is recommended for gBLUP, and GAPIT is preferred over other methods for using cBLUP and sBLUP.

**Figure 6.**
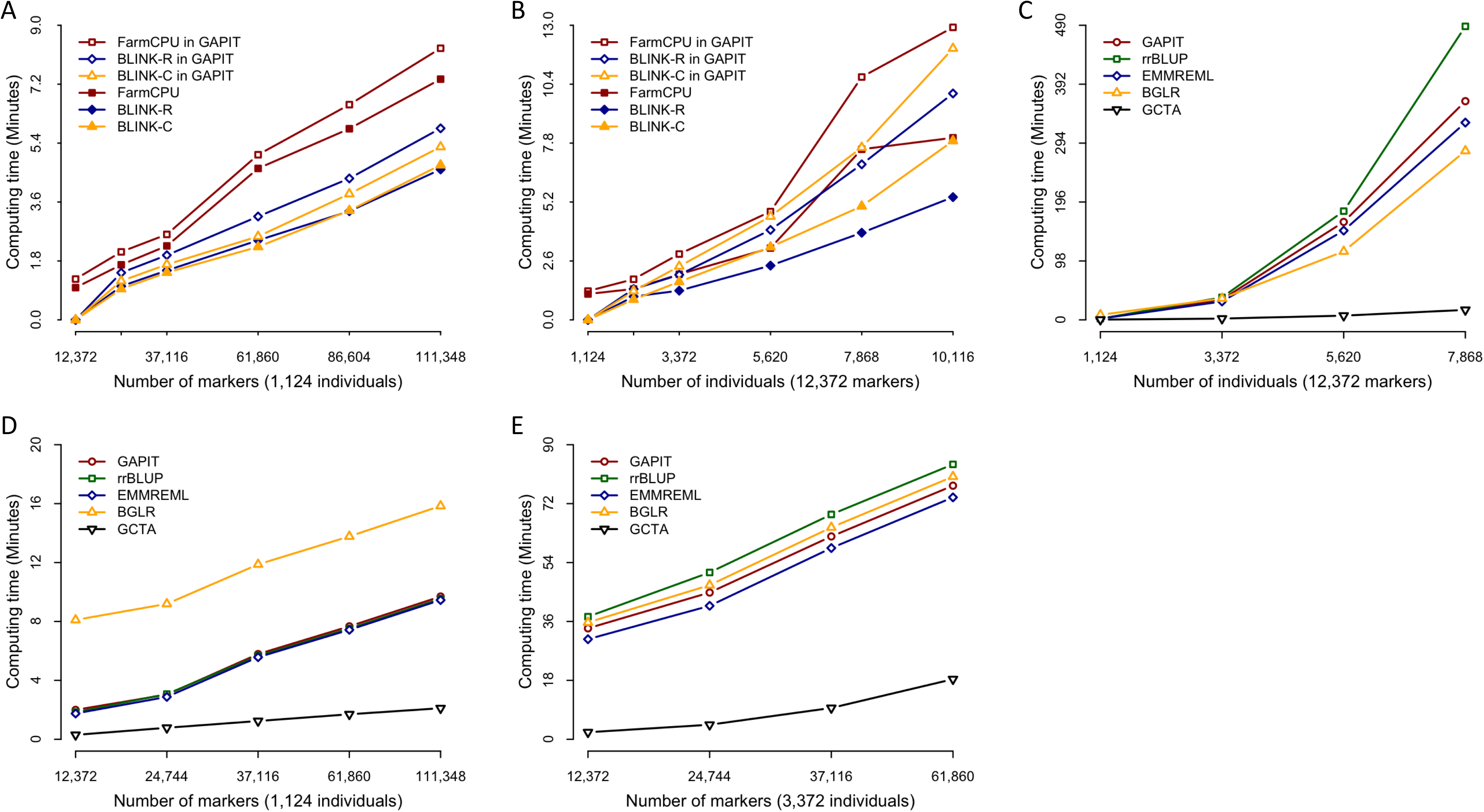
Comparison of computing time using multiple packages of GWAS and GS within and outside of GAPIT. Three GWAS packages (FarmCPU, BLINK C version and BLINK R version) were compared by running them within GAPIT and outside of GAPIT as standalone. The comparison was conducted on a synthetic trait simulated from the maize genotypes (281 individuals and 3093 markers). The trait was simulated with 75% heritability controlled by 20 QTNs. To demonstrate the impact on computing time, the data was duplicated for markers (A) and individuals (B) at multiple times (8, 12, 20, 28, and 36). Either running within GAPT or outside of GAPIT as standalone, these GWAS packages exhibit linear computing time to both number of markers and number of individuals. The extra time of execution of these packages within GAPIT is minimal comparing to the execution as standard alone. The extra time involves format transformation of input date and result presentation. Computing time was compared for five packages of genomic prediction, including GAPIT, GCTA, BGLR, rrBLUP, and EMMAREML. The genomic Best Linear Unbiased Prediction was selected in GAPIT. With number of markers duplicated four times and number of individuals duplicated at multiple levels (12, 20, and 28 fold), the computing show nonlinear relationship to number of individuals, except the GCTA package (C). With number of individual duplicated 4 (D) and 12 (E) times; and number of markers duplicated at multiple levels (12, 20, 28, and 36 fold), the computing time show linear relationship to number of marker for all package. The numbers of individuals change the rank of the packages. BGLR is the slowest with less individuals (D) and rrBLUP become the slowest with more individuals (E).

## Discussion

### Comprehensive and specific software packages

Developments of sophisticated and computationally efficient methods are essential for genomic research. Software initiation, upgrade, and maintenance are equally crucial for turning genomic data into knowledge. These software packages can be classified into two categories: specific and comprehensive. Packages in the specific category are usually accompanied by the development of new methods, such as MLMM[14], FarmCPU[15], and BLINK[16]. Due to the limitation of time and resources, these software packages target the implementation of specific methods with a direct link between input data and output, mainly the p values. This type of software package does not provide comprehensive functions for input data diagnosis or output results interpretation. Consequently, users must rely on other types of software packages (comprehensive) to complete their analyses.

Some software packages may initiate as a specific package, but build functions over time to become comprehensive. One example is TASSEL. Alternatively, some software packages, such as PLINK[23], BGLR [29], rrBLUP[25], GCTA[30], iPAT[31], and GAPIT[27,28], are designed to be comprehensive from the start. Originally, GAPIT1 implemented GLM, MLM, and CMLM for GWAS and gBLUP for GS. GAPIT1 also provided a comprehensive report, including many figures and tables that can be used in publications. In GAPIT2, we added four new methods for GWAS, including FaST-LMM, FaST-LMM-Select, ECMLM, and SUPER, and updated the report outputs. In the current GAPIT3, we added three multiple locus test methods for GWAS (MLMM, FarmCPU, and BLINK) and two methods for GS (cBLUP and sBLUP).

The learning curves for the two types of software packages, specific and comprehensive, vary across users and packages. Some users are eager to learn new software packages, especially the specific software packages that are more straightforward. In contrast, some users are comfortable with their existing knowledge and skills, especially when they have mastered a particular comprehensive software package. GAPIT3 targets both types of users. For users that are new to GAPIT, we designed simple prompts and commands: “tell me your genotype and phenotype data, we do our best.” For existing users, we maximized the consistency between versions such as typing commands, selecting options, and navigating reports and graphics to obtain information. For example, to choose a GWAS method among the ten available methods in GAPIT3, users simply add the model statement as in previous GAPIT versions. According to the GAPIT forum, no difficulties have been expressed in using GAPIT3 compared to previous versions.

### Selection of GWAS and GS methods

Although the current architecture of GAPIT3 makes is easy to implement an R package, selection of methods is critical for boosting statistical power and accuracy for GWAS and GS. We used the gaps of implementations and performance as the criteria for the selection of these packages. The method of fitting all markers simultaneously as random effects as an alternative to gBLUP for GS was introduced in 2001 [32]. The ridge regression and Bayes theory-based methods (e.g., Bayes A, B, and CPi) can be used not only to predict individuals’ breeding values by summing the effects of all markers, but also to map genetic markers associated with phenotypes of interest [33]. Multiple comprehensive software packages have been developed for both GWAS and GS, including BGLR [29], rrBLUP [21], GCTA [30].

For the conventional method of single-locus test, many advanced methods were developed, including incorporation of population structure [2], kinship [34], compressed kinship [35], and complementary kinship [11,36]. Many software packages were developed for these specific methods, including EMMA, EMMAx, FaSTLMM, GEMMA, and GenABEL. Comprehensive software packages, including PLINK, TASSEL, and GAPIT, were also developed to implement many of these methods.

The multiple-locus test, evolved over time to use the format of stepwise regression with a fixed effect model, for example, the SAS GLMSELECT procedure [37], or with a mixed model, for example, the R package of MLMM [38]. Furthermore, the stepwise regression format was advanced to the iteration of two models. The first model is used to test markers one at a time, and the second model is used to evaluate the associated markers as cofactors in the first model to re-test markers [15,16]. Two different iterative models are available: FarmCPU and BLINK. FarmCPU uses a fixed effect model and a random effect model. BLINK uses two fixed effect models. Related studies have demonstrated that multiple-locus methods are generally superior to single-locus methods. With the exception of GLMSELECT by SAS, multiple-locus methods for GWAS have yet to be implemented in a comprehensive software package[39]. Consequently, we chose to implement FarmCPU and BLINK in GAPIT3 to boost statistical power for GWAS.

For GS, GAPIT1 implemented gBLUP, which is superior for traits controlled by a large number of genes, but not as effective for traits controlled by a small number of genes. In GAPIT3, we implemented a newly developed method, sBLUP, which is superior to gBLUP for such traits. The common problem for both gBLUP and sBLUP is their lack of effectiveness when executing GS for traits with low heritability. Therefore, in the updated GAPIT3, we implemented a newly developed method, cBLUP, which is superior for traits with low heritability. By doing so, GAPIT3 performs well across the full spectrum of traits, whether controlled by a large or small number of genes and with either high or low heritability.

### Operation of GAPIT

GAPIT is an R package executed through the command-line interface (CLI), which is efficient for repetitive analyses such as multiple traits and using multiple methods and models. However, CLI is not as straightforward as the software packages equipped with a graphical user interface (GUI), such as TASSEL and iPAT. Instead, GAPIT requires users to input some keywords in specific formats. The advantage of living in the age of the Internet, is that we can transform peoples’ excellent reading, copying, and pasting skills into actions that reduce the complexities of executing GAPIT. We provide ~20 tutorials on the GAPIT website that users can read, edit, copy, and paste as necessary to efficiently use the CLI to conduct most of the analyses.

### Limitations

As an R package, GAPIT faces challenges when dealing with big data. Most of the analyses using GAPIT require data to be loaded into memory. However, the FarmCPU can use a R package (bigmemory) to import big data and carry all analyses into the final P values. The current GAPIT team is currently working on this feature. For users with big data, a viable option is to run GAPIT with the BLINK C version, which only reads data pertinent to the analyses from a specific section on the disk/drive. The only requirement is an executable file of the BLINK C version in the working directory of R.

## Conclusion

GAPIT has served the genomic research community for eight years, since 2012, as a Genomic Association and Prediction Tool in the form of an R package. The software is extensively used worldwide, as indicated by over 800 citations of two publications (Bioinformatics in 2012 and The Plant Genome in 2016), ~2000 posts on GAPIT forum, and ~22,000 page views on the GAPIT website. In the new GAPIT3, we implemented three multiple-loci test methods (MLMM, FarmCPU, and BLINK) for GWAS and two more variations of BLUP (compressed BLUP and SUPER BLUP) for genomic selection. GAPIT3 also includes enhancements to the analytical reports as part of our continuous efforts to build upon the comprehensive output reports developed in versions 1 and 2. These enhancements assist users in the interpretation of input data and analytical results. Valuable new features include the users’ ability to instantly and interactively extract information for individuals and markers on Manhattan plots, QQ plots, and scatter plots of predicted vs. observed phenotypes.

## Supporting information

https://www.researchgate.net/publication/346482922_GAPIT3_Supplementary

https://www.researchgate.net/publication/346482682_Table1

## Availability

The GAPIT source code, demo script, and demo data are freely available on the GAPIT website (www.zzlab.net/GAPIT).

### Acknowledgment

The authors thank Linda R. Klein for helpful comments and editing the manuscript. This project was partially funded by National Science Foundation, the United States (Award # DBI 1661348 and ISO 2029933), the United States Department of Agriculture-National Institute of Food and Agriculture, the United States (Hatch project 1014919, Award #s 2016-68004-24770, 2018-70005-28792, and 2019-67013-29171), the Washington Grain Commission, the United States(Endowment and Award #s 126593 and 134574), the Program of Chinese National Beef Cattle and Yak Industrial Technology System, China (Award #s CARS-37), Fundamental Research Funds for the Central Universities, China (Southwest Minzu University, Award #s 2020NQN26), and Sichuan Science and Technology Program, China (Hatch project 21YYJC2934 and 21YYJC2967).

## Supplementary material

**Figure S1.**
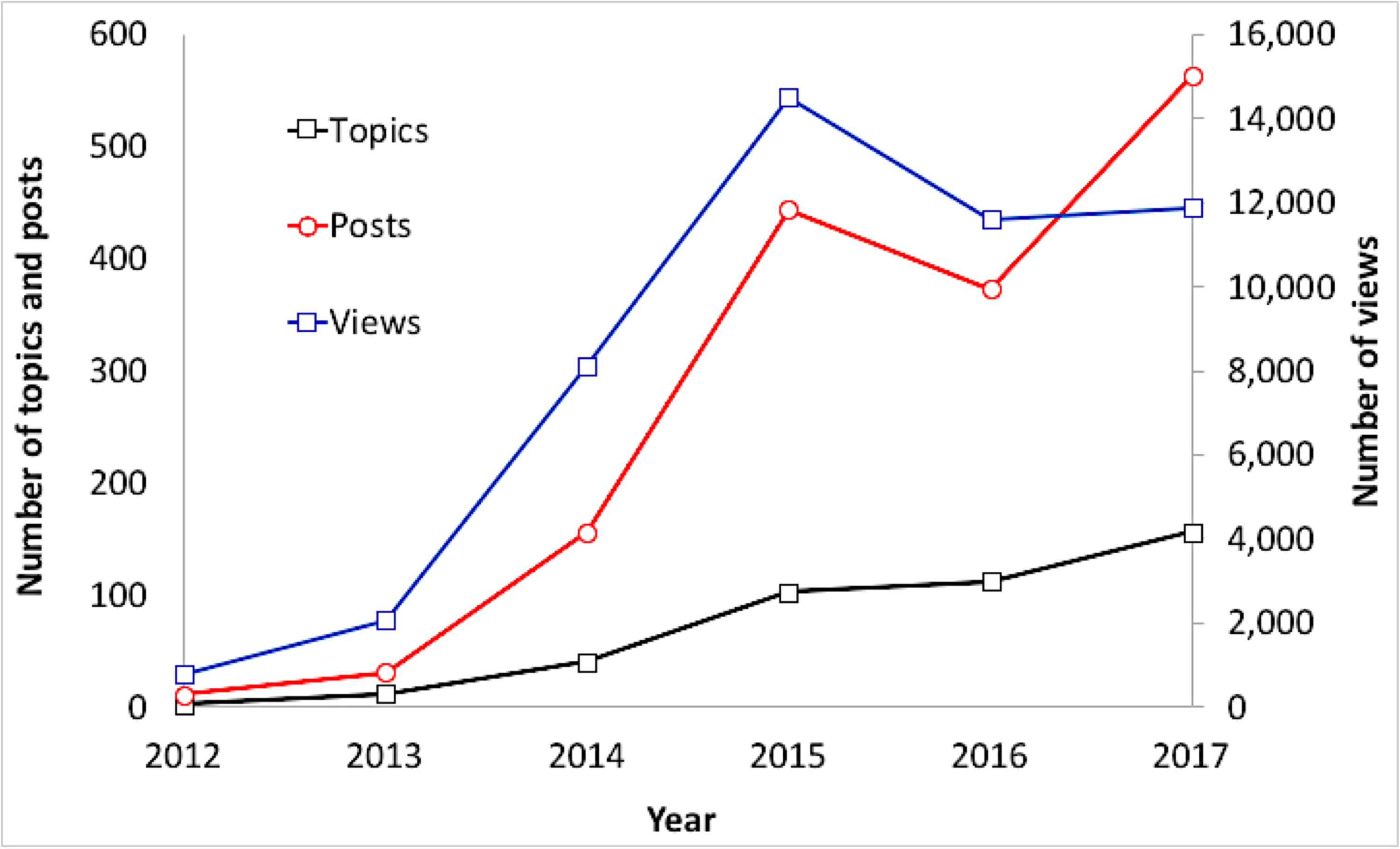
Interaction among users and developers on GAPIT forum through Google. Since the first post in 2012, the forum has received over 700 topics, 3,000 posts and 80,000 views in total. This trend is increasing overall for all three measurements. Exceptions were observed in 2016 and 2019, corresponding to the 2016 event when Google was withheld from users in China and the restriction of accessing Google using VPN (https://en.wikipedia.org/wiki/Google_China).

**Figure S2.**
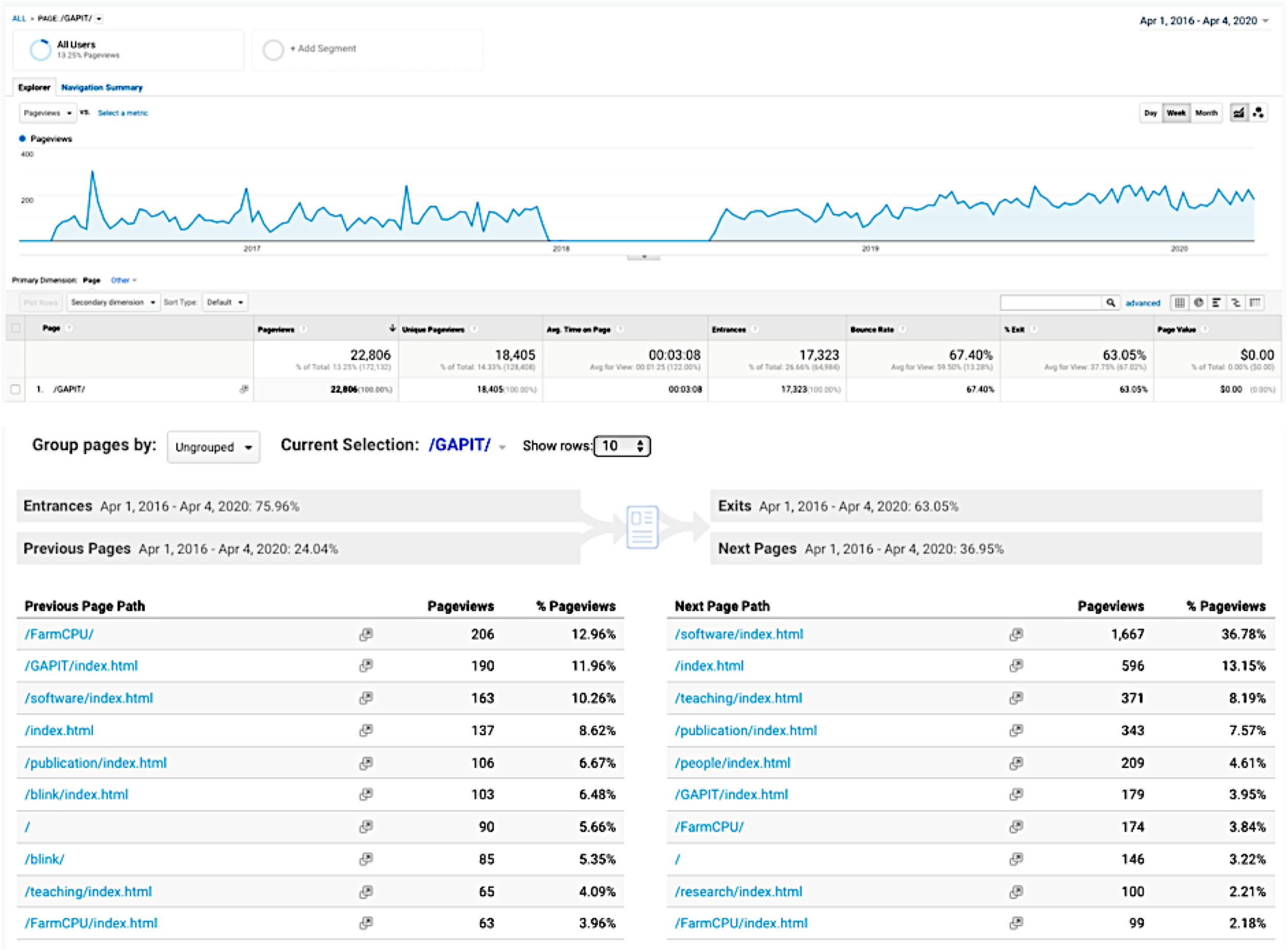
Usage of GAPIT website. The GAPIT website has received 22,806 page views since 2016 when we began tracking the usage on Google Analytics. We lost about six months of tracking due to a technology issue. The average page view time is three minutes and eight seconds, accounting for 49.6 days in total. An increasing trend for weekly total number of page views is observed, which is currently over 200 pageviews per week. The previous page paths are FarmCPU (17%), BLINK (12%), Publication (7%), and teaching (4%). The majority of next page paths are software pages, which host several software packages developed at Zhiwu Zhang Lab, including FarmCPU and BLINK for GWAS, and GRID and GridFree for image analyses.

**Figure S3.**
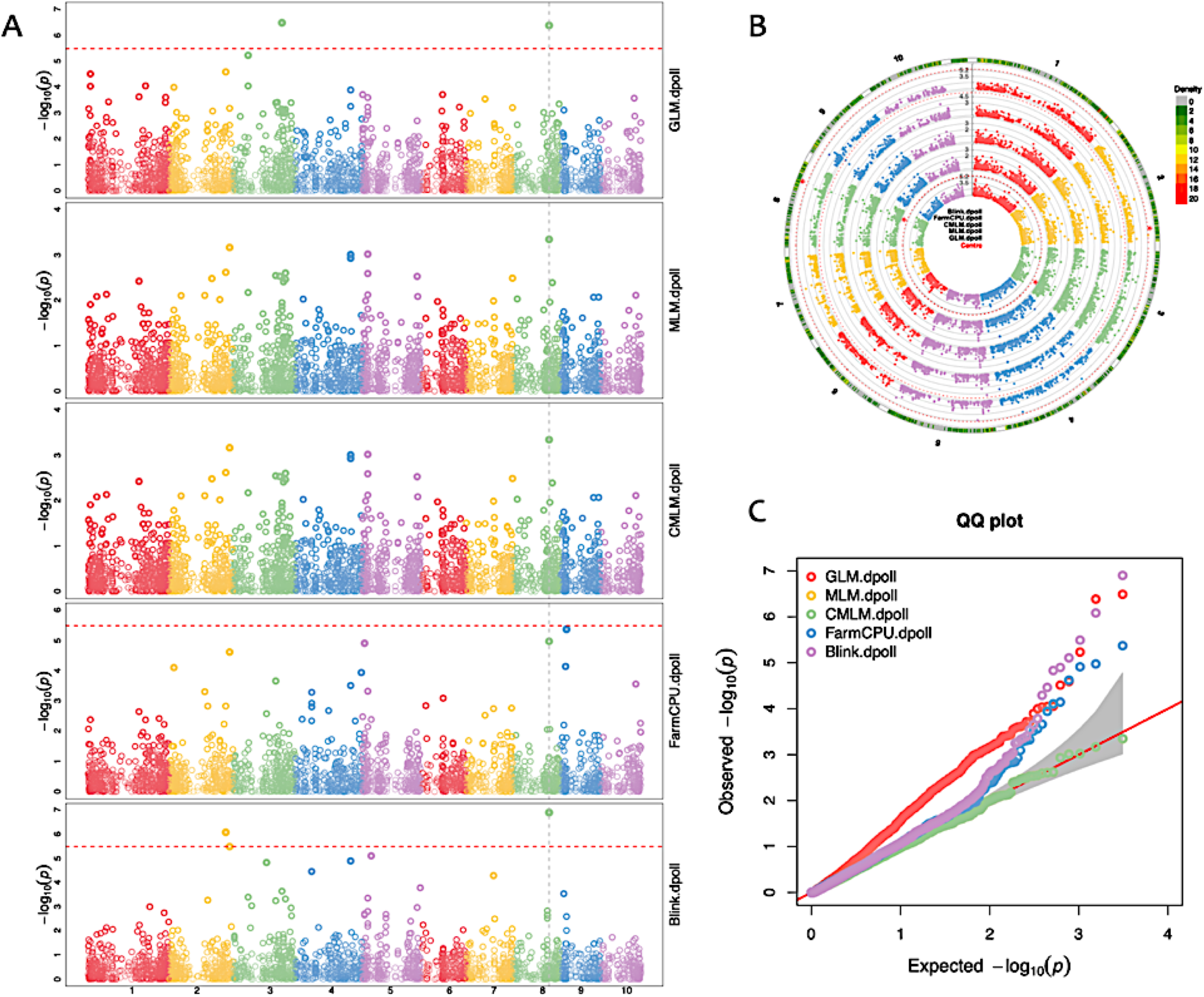
Interactive Manhattan and QQ plots. As a software package that includes multiple GWAS methods, GAPIT supplies the user with interactive Manhattan and QQ plots to compare results among the methods selected. Two types of Manhattan plots are displayed, the standard orthogonal plot (A) and a circle plot (B). A multiple method QQ plot is also displayed (C). Each method’s Manhattan plot includes an interconnected, dashed vertical line that runs through chromosome 8, signaling that only two methods have detected this association signal (i.e., potentially significant SNP) with the peak p-value. In contrast, a solid (not dashed) vertical line is displayed if more than two methods detect the same signal with the peak p-value. The circle plot also supplies a marker distribution analysis, represented by the colors, ranging from green to red, in the outermost ring. Areas in the outer ring that are colored red have the greatest number of markers within the selected window size (10Kbp is the default, but can be changed by the user).

**Figure S4.**
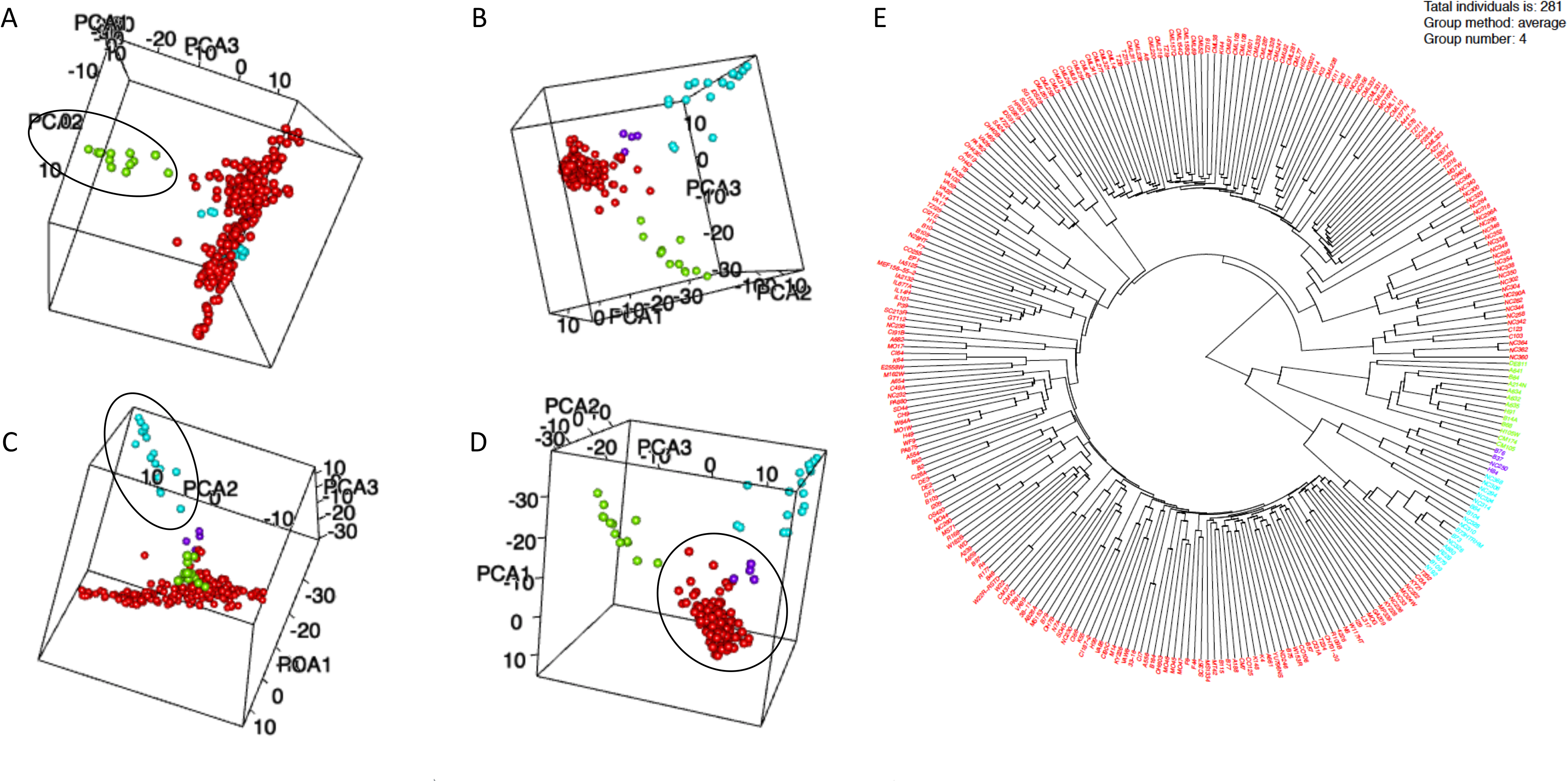
Interactive display of population structure and kinship cladogram. Population structure is displayed as an interactive three-dimension plot. Users can adjust the display at any angle (e.g., A to D). The individuals are displayed with colors that correspond to the grouping on the kinship cladogram using k-means cluster analysis (E).

**Figure S5.**
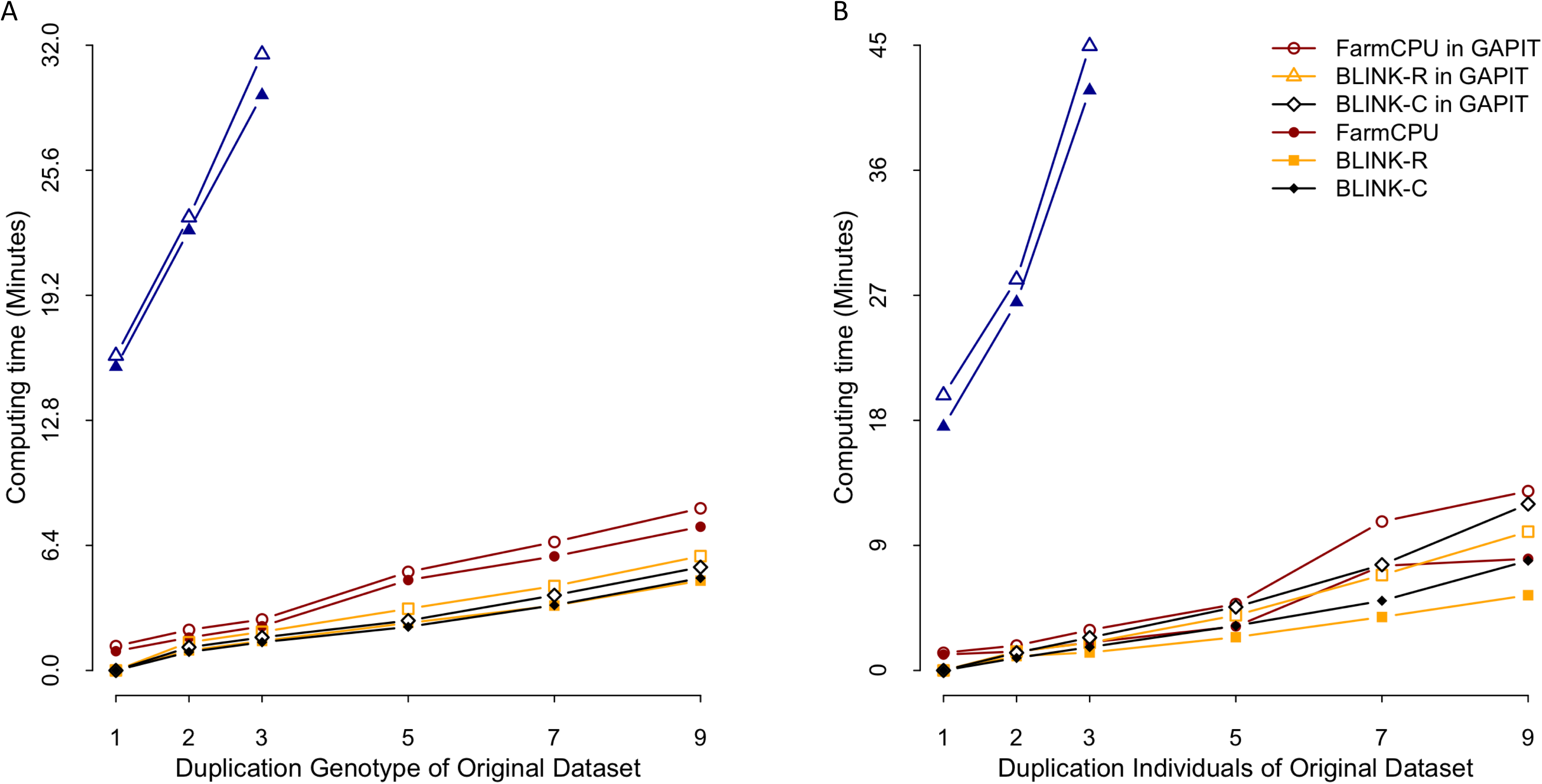
Comparison of computing time using four software packages run separately and using them within GAPIT. The three standalone software packages are MLMM, FarmCPU, BLINK R version, and BLINK C version. The comparison was performed on different sized datasets with respect to duplication of the original data containing 1124 individuals and 12,372 markers. The duplications were conducted for markers only (A) and individuals only (B). In either case, these packages exhibit linear computing time to number of markers, and number of individuals. The extra time of execution of these packages within GAPIT is minimal comparing to the execution as standard alone. The extra time involves format transformation of input date and result presentation. MLMM took much longer time than the rest three packages, which are not able to be differentiated each other when they displayed on the same scale with MLMM.

**Figure S6.**
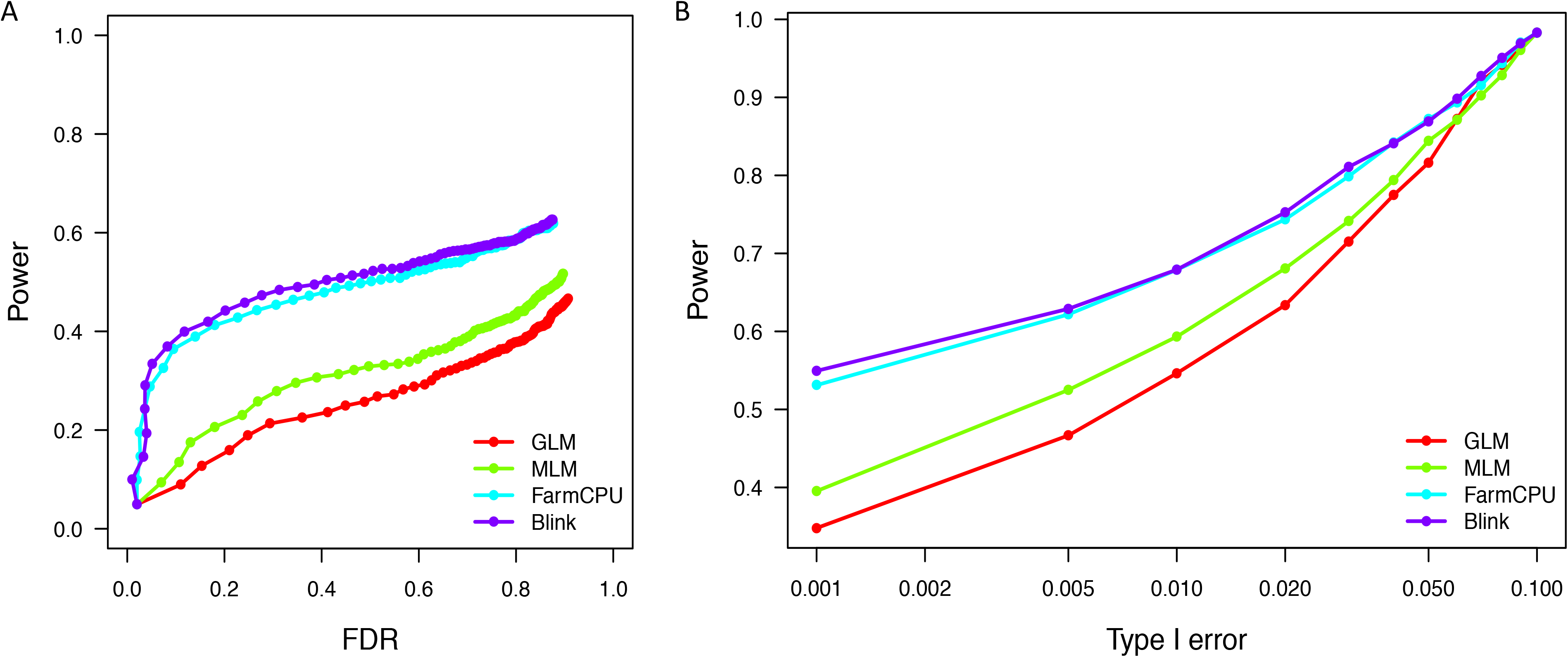
Comparison between single locus and multiple loci methods on power against FDR and Type I error. Single-locus methods include GLM and MLM. The Multi-loci methods include FarmCPU and Blink. The comparison was based a simulated trait using the maize data containing 282 individuals and 3094 SNPs. The simulated trait had a heritability of 75% controlled by 20 Quantitative Trait Nucleotides (QTN). Power was calculated as the proportion of QTN detected. False Discover Rate (FDR) was calculated as the proportion of non-QTNs among the positives (A). Type I error was calculated as the proportion of tests with false positives (B). The simulation was replicated 100 times.

