## Supplementary material for "GAPIT Version 3: Boosting Power and Accuracy for Genomic Association and Prediction": https://www.researchgate.net/publication/346482922_GAPIT3_Supplementary

**Supplementary documents**

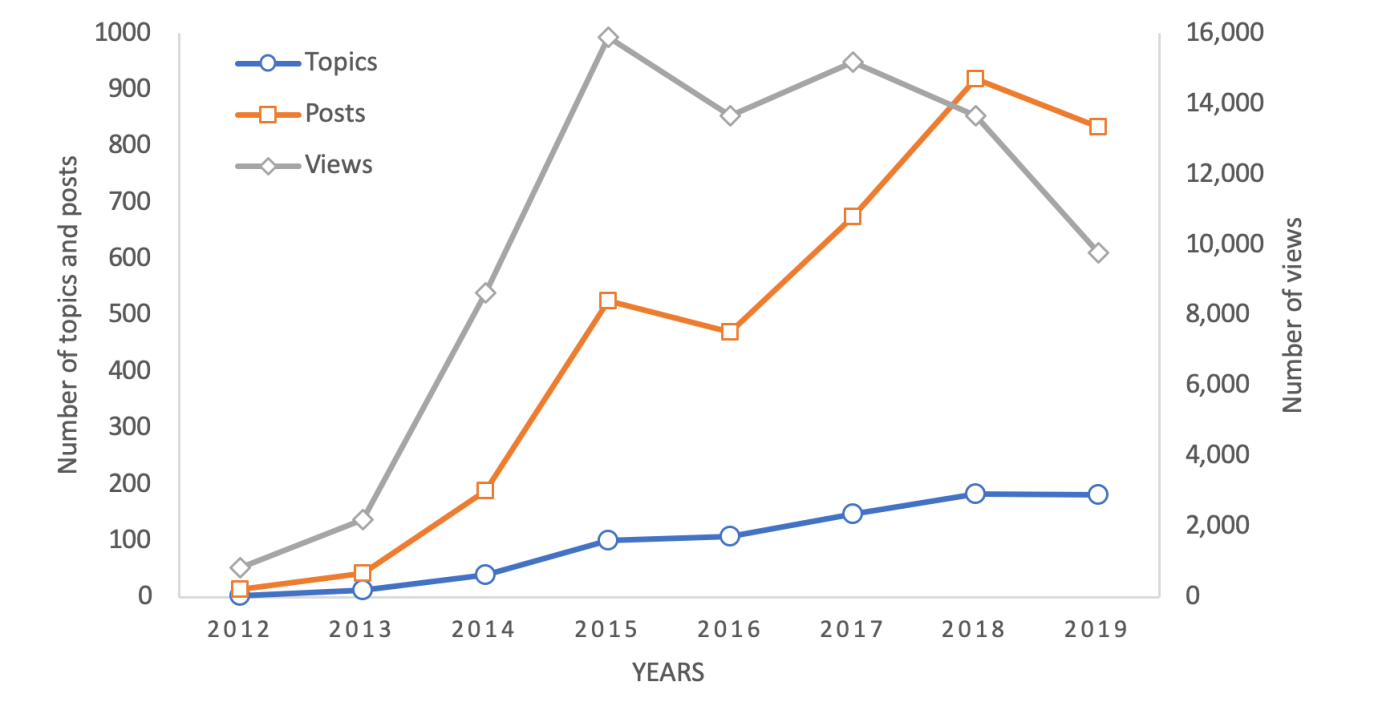

**Figure S1. Interaction among users and developers on GAPIT forum through Google.** Since the first post in 2012, the forum has received over 700 topics, 3,000 posts and 80,000 views in total. This trend is increasing overall for all three measurements. Exceptions were observed in 2016 and 2019, corresponding to the 2016 event when Google was withheld from users in China and the restriction of accessing Google using VPN (<https://en.wikipedia.org/wiki/Google_China>).

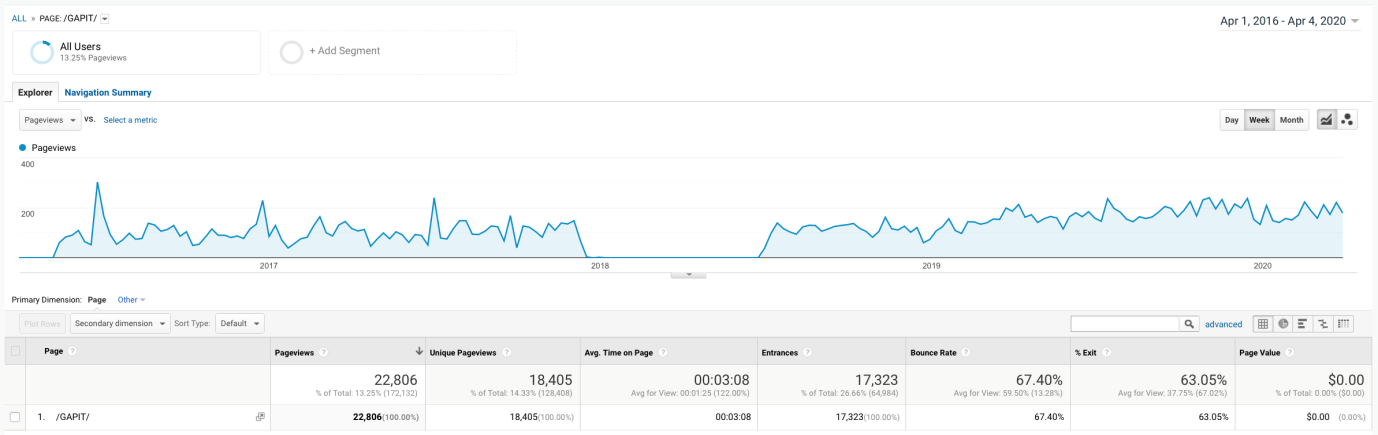

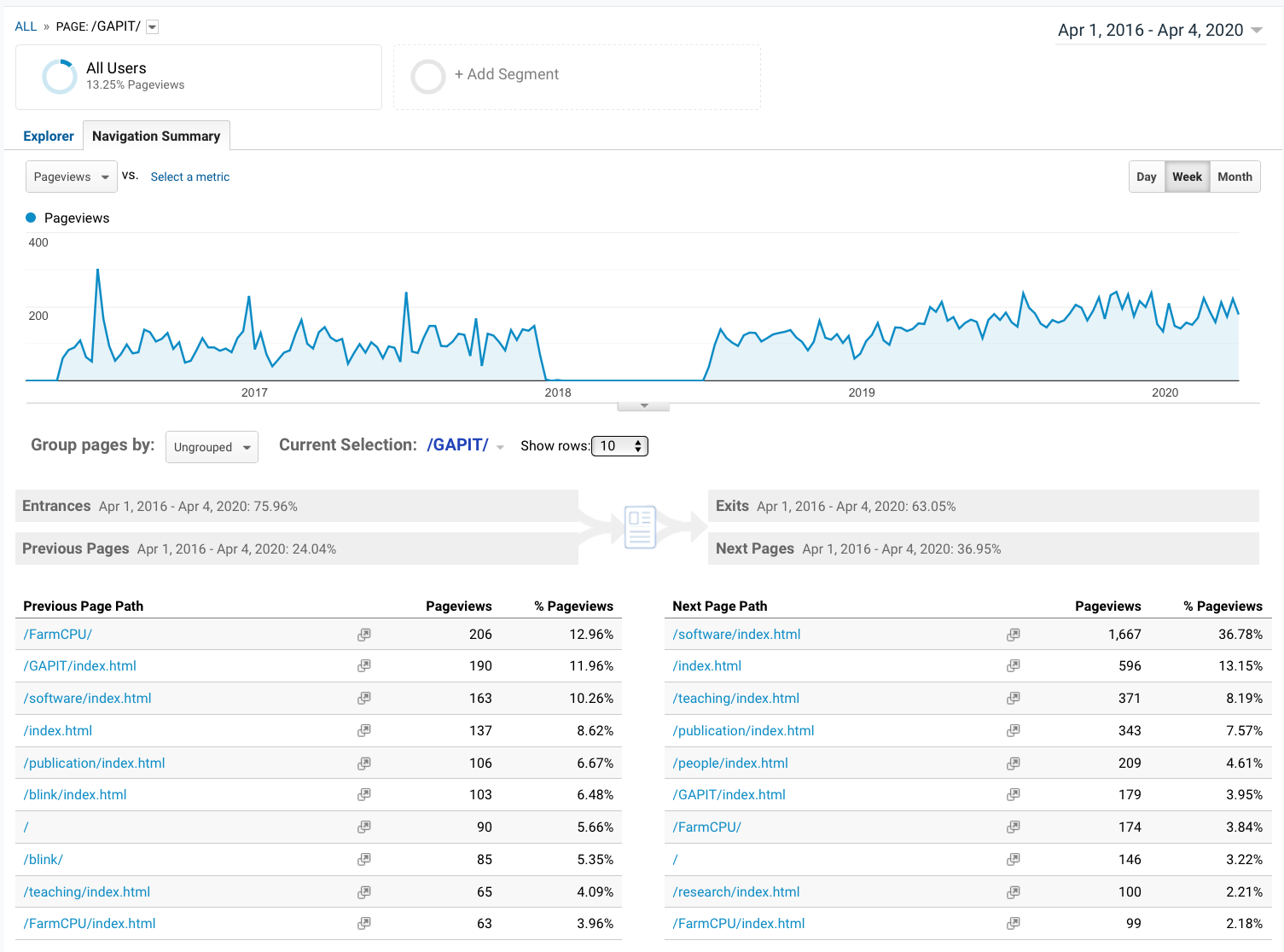

**Figure S2. Usage of GAPIT website.** The GAPIT website has received 22,806 page views since 2016 when we began tracking the usage on Google Analytics. We lost about six months of tracking due to a technology issue. The average page view time is three minutes and eight seconds, accounting for 49.6 days in total. An increasing trend for weekly total number of page views is observed, which is currently over 200 pageviews per week. The previous page paths are FarmCPU (17%), BLINK (12%), Publication (7%), and teaching (4%). The majority of next page paths are software pages, which host several software packages developed at Zhiwu Zhang Lab, including FarmCPU and BLINK for GWAS, and GRID and GridFree for image analyses.

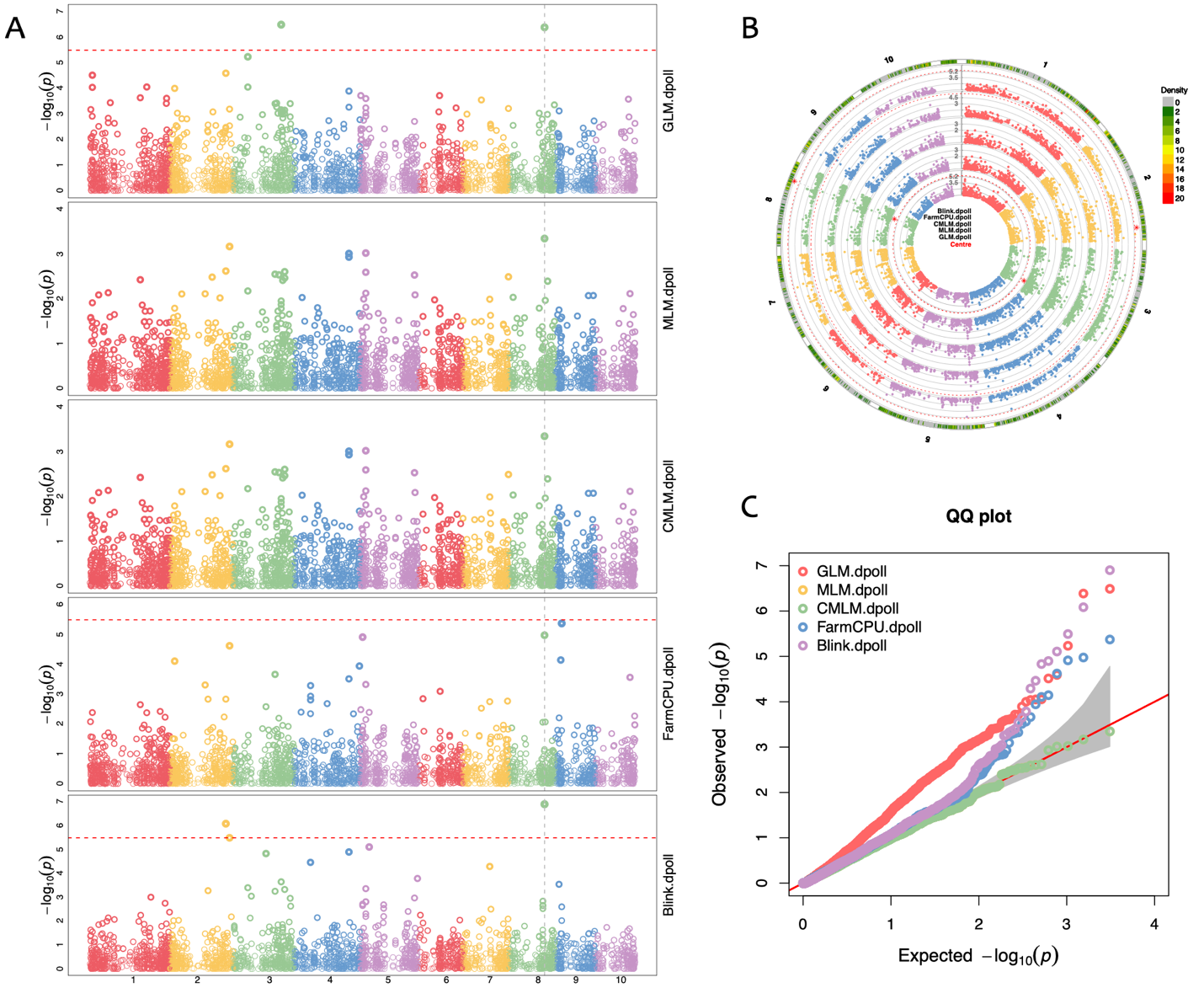

**Figure S3. Interactive Manhattan and QQ plots.** As a software package that includes multiple GWAS methods, GAPIT supplies the user with interactive Manhattan and QQ plots to compare results among the methods selected. Two types of Manhattan plots are displayed, the standard orthogonal plot (A) and a circle plot (B). A multiple method QQ plot is also displayed (C). Each method's Manhattan plot includes an interconnected, dashed vertical line that runs through chromosome 8, signaling that only two methods have detected this association signal (i.e., potentially significant SNP) with the peak p-value. In contrast, a solid (not dashed) vertical line is displayed if more than two methods detect the same signal with the peak p-value. The circle plot also supplies a marker distribution analysis, represented by the colors, ranging from green to red, in the outermost ring. Areas in the outer ring that are colored red have the greatest number of markers within the selected window size (10Kbp is the default, but can be changed by the user).

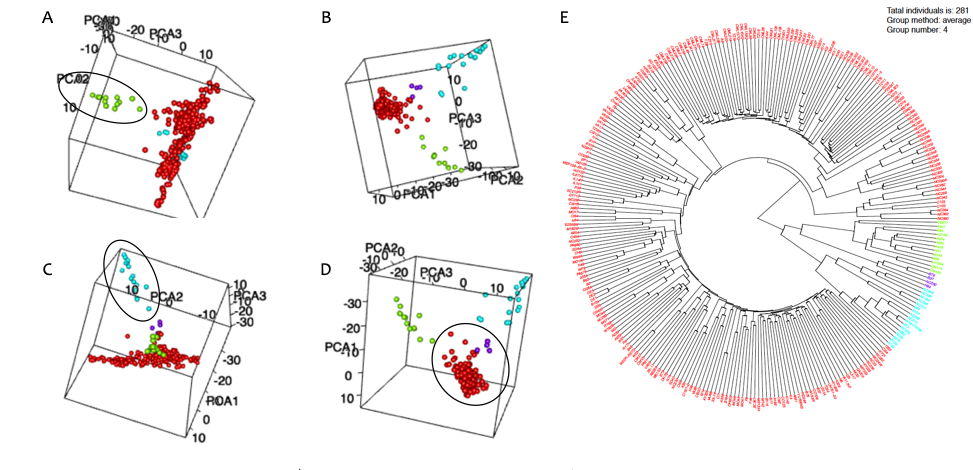

**Figure S4. Interactive display of population structure and kinship cladogram.**

Population structure is displayed as an interactive three-dimension plot. Users can adjust the display at any angle (e.g., A to D). The individuals are displayed with colors that correspond to the grouping on the kinship cladogram using k-means cluster analysis (E).

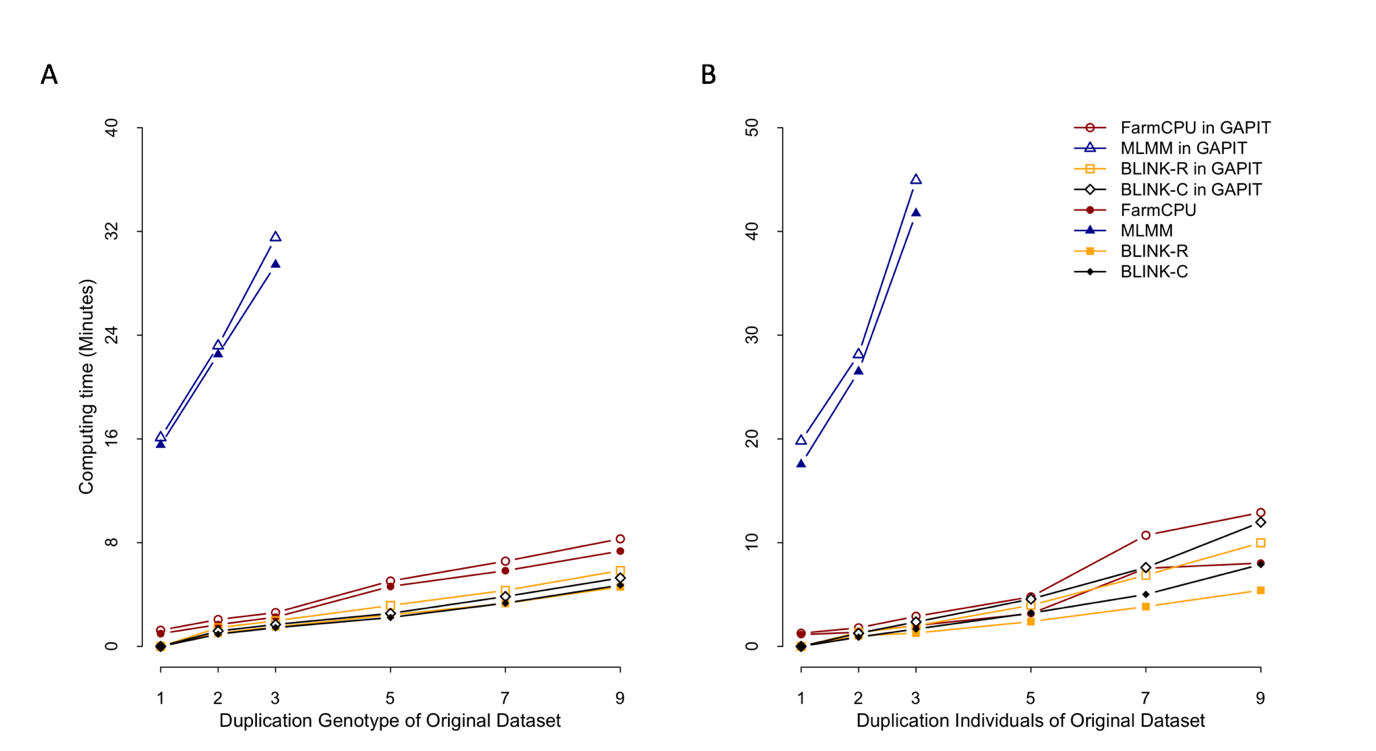

**Figure S5. Comparison of computing time using four software packages run separately and using them within GAPIT.** The three standalone software packages are MLMM, FarmCPU, BLINK R version, and BLINK C version. The comparison was performed on different sized datasets with respect to duplication of the original data containing 1124 individuals and 12,372 markers. The duplications were conducted for markers only (A) and individuals only (B). In either case, these packages exhibit linear computing time to number of markers, and number of individuals. The extra time of execution of these packages within GAPIT is minimal comparing to the execution as standard alone. The extra time involves format transformation of input date and result presentation. MLMM took much longer time than the rest three packages, which are not able to be differentiated each other when they displayed on the same scale with MLMM.

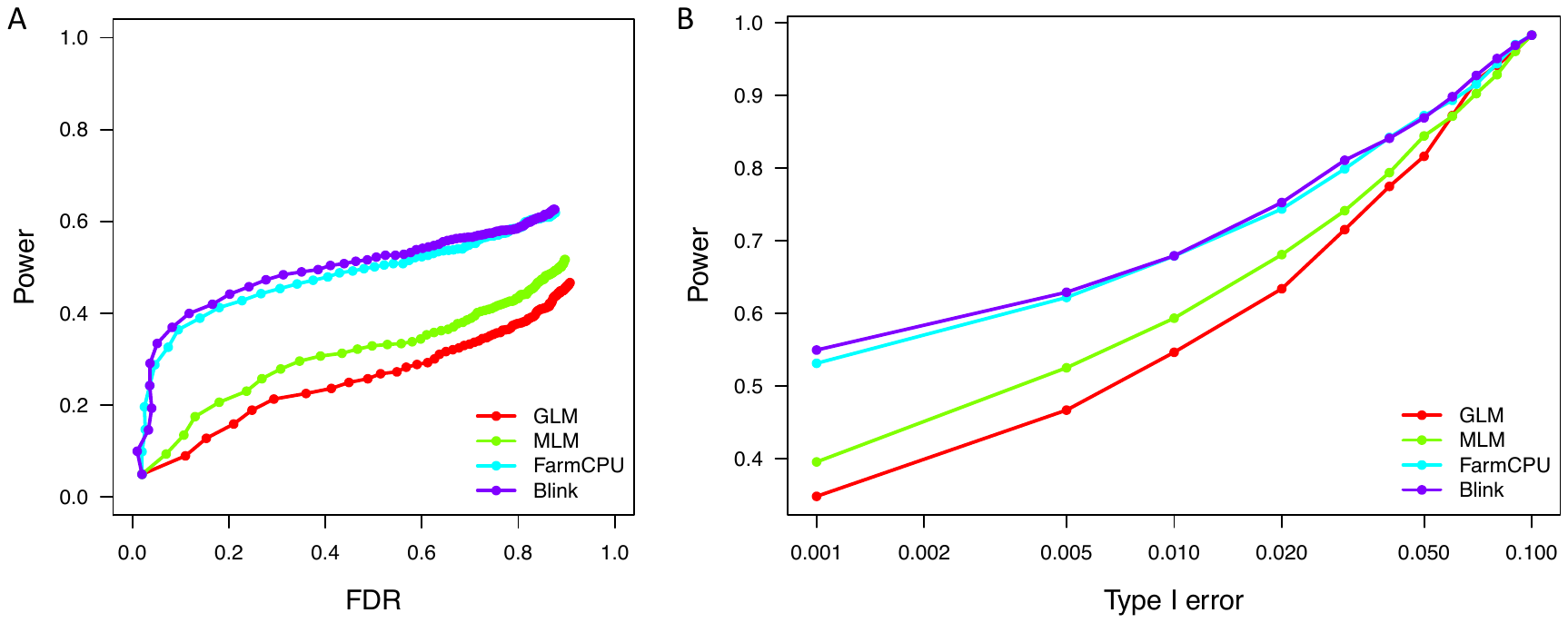

**Figure S6. Comparison between single locus and multiple loci methods on power against FDR and Type I error.** Single-locus methods include GLM and MLM. The Multi-loci methods include FarmCPU and Blink. The comparison was based a simulated trait using the maize data containing 282 individuals and 3094 SNPs. The simulated trait had a heritability of 75% controlled by 20 Quantitative Trait Nucleotides (QTN). Power was calculated as the proportion of QTN detected. False Discover Rate (FDR) was calculated as the proportion of non-QTNs among the positives (A). Type I error was calculated as the proportion of tests with false positives (B). The simulation was replicated 100 times.
