## Supplementary material for "GAPIT Version 3: Boosting Power and Accuracy for Genomic Association and Prediction": https://www.researchgate.net/publication/346482682_Table1

**Table 1. Characteristics of methods in GAPIT3.**

| **Method** | **Testing Marker** | **Steps** | **Model** | **Kinship** |
| --- | --- | --- | --- | --- |
| GLM | Single-locus | one | Fixed | NA |
| MLM | Single-locus | one | Mixed | All markers |
| CMLM | Single-locus | one | Mixed | Cluster individuals into groups |
| ECMLM | Single-locus | one | Mixed | Enrichment cluster individuals into groups |
| SUPER | Single-locus | two | Mixed | All marker except pseudo QTNs |
| MLMM | Multiple-loci | Iterative | Mixed | All markers |
| FarmCPU | Multiple-loci | Iterative | Fixed and Mixed | Pseudo QTNs |
| BLINK | Multiple-loci | Iterative | Fixed | NA |
| gBLUP | NA | one | Mixed | All markers for all individuals |
| cBLUP | NA | one | Mixed | Cluster individuals into groups with all markers |
| sBLUP | NA | one | Mixed | Pseudo QTNs |
